# The Rous sarcoma virus Gag polyprotein forms biomolecular condensates driven by intrinsically-disordered regions

**DOI:** 10.1101/2023.04.07.536043

**Authors:** Rebecca Kaddis Maldonado, Breanna L. Rice, Gregory S. Lambert, Malgorzata Sudol, John M. Flanagan, Leslie J. Parent

## Abstract

Biomolecular condensates (BMCs) play important roles in cellular structures including transcription factories, splicing speckles, and nucleoli. BMCs bring together proteins and other macromolecules, selectively concentrating them so that specific reactions can occur without interference from the surrounding environment. BMCs are often made up of proteins that contain intrinsically disordered regions (IDRs), form phase-separated spherical puncta, form liquid-like droplets that undergo fusion and fission, contain molecules that are mobile, and are disrupted with phase-dissolving drugs such as 1,6-hexanediol. In addition to cellular proteins, many viruses, including influenza A, SARS-CoV-2, and human immunodeficiency virus type 1 (HIV-1) encode proteins that undergo phase separation and rely on BMC formation for replication. In prior studies of the retrovirus Rous sarcoma virus (RSV), we observed that the Gag protein forms discrete spherical puncta in the nucleus, cytoplasm, and at the plasma membrane that co-localize with viral RNA and host factors, raising the possibility that RSV Gag forms BMCs that participate in the virion intracellular assembly pathway. In our current studies, we found that Gag contains IDRs in the N-terminal (MAp2p10) and C-terminal (NC) regions of the protein and fulfills many criteria of BMCs. Although the role of BMC formation in RSV assembly requires further study, our results suggest the biophysical properties of condensates are required for the formation of Gag complexes in the nucleus and the cohesion of these complexes as they traffic through the nuclear pore, into the cytoplasm, and to the plasma membrane, where the final assembly and release of virus particles occurs.

## Introduction

Rous sarcoma virus (RSV), an avian oncoretrovirus discovered by Peyton Rous in 1910 [1–4], was the first virus found to cause solid tumors [1, 5, 6]. The RSV Gag polyprotein, which orchestrates the assembly of nascent virions, has served for decades as the basis for dissecting the modular assembly domains involved in virus particle biogenesis [7–14]. Gag is synthesized as a multidomain precursor that is proteolytically cleaved after budding by the C-terminal protease (PR) domain into the mature matrix (MA), p2, p10, capsid (CA), and nucleocapsid (NC) that make up the virion core (reviewed in [15]). Based on the observation that virus particles are released from the plasma membrane, it was generally accepted that RSV Gag functioned exclusively in the cytoplasm. However, using genetic, biochemical, and advanced imaging approaches, we discovered that RSV Gag undergoes transient nucleocytoplasmic trafficking. This process is mediated by intrinsic nuclear localization signals (NLS) in MA and NC, and a nuclear export signal (NES) in p10 that interacts with the CRM1-RanGTP export complex to exit through the nuclear envelope [16–23].

To examine the mechanisms underlying RSV Gag subcellular trafficking, we have studied the localization of wild-type and mutant Gag proteins fused to fluorescent tags [16-19, 22-24], in both living and fixed cells. These imaging experiments revealed that Gag forms discrete foci in the nucleus, cytoplasm, and at the plasma membrane ([4]; Fig 1). Electron micrographs have failed to demonstrate that there are membranes surrounding these intracellular Gag foci (unpublished data), raising the possibility that Gag forms well-defined, focal puncta using similar mechanisms to those that govern the formation of membraneless organelles--also known as biomolecular condensates (BMCs) [3, 22, 25–34]—in nuclear and cytoplasmic compartments, which accumulate along the plasma membrane for release from the cell. ^1^

**Figure 1:**
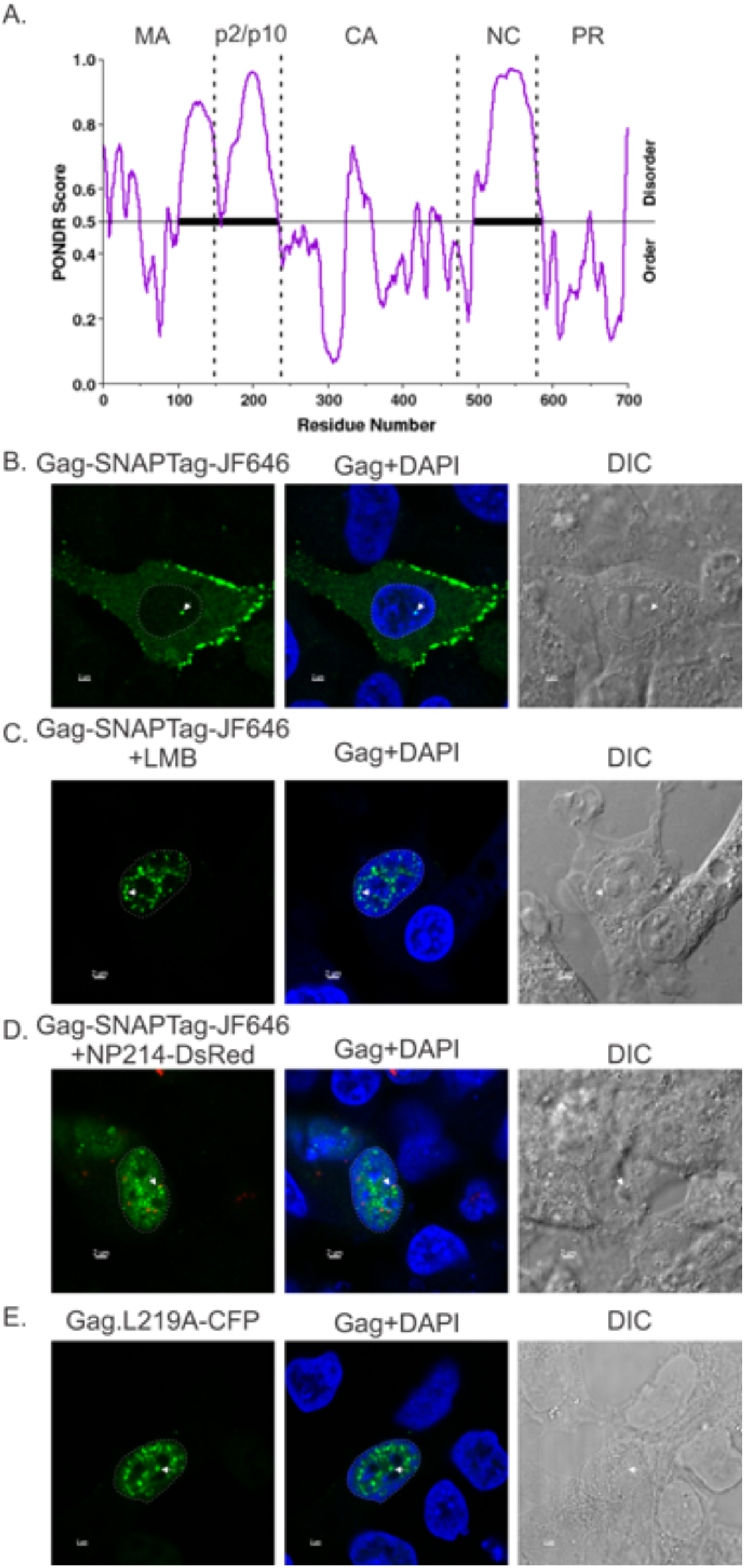
RSV Gag contains intrinsically disordered regions (IDRs) and forms BMCs in cells. (A) PONDR analysis of RSV reveals two IDRs, one spanning Map2p10 and the other within NC. (B) Wild-type Gag-SNAPTag labeled with JF646 SNAP ligand (green) formed foci along the plasma membrane, in the cytoplasm, and in the nucleus (DAPI, blue; nucleus, white outline). The same phenomenon can be viewed when WT Gag-SNAP-tag JF646 was trapped in the nucleus either by (C) CRM1 inhibition by leptomycin B (LMB) or (D) a dominant-negative of Nup214 (NP214-DsRed). (E) The Gag.L219A-CFP (green) nuclear export mutant formed phase-contrasted nuclear foci that can be viewed using DIC, a known characteristic of BMCs. Scale bar = 2 µm.

To organize cellular activities into discrete compartments that are not membrane-enclosed, macromolecules can form densely packed, discrete complexes known as BMCs [34, 35]. As the protein concentration in these complexes reaches a critical threshold, de-mixing occurs and liquid-like droplets form, physically separating themselves from the surrounding milieu and adopting properties of a liquid [27, 34–37]. Phase separation is driven by multivalent protein-protein and protein-nucleic acid interactions consisting of electrostatic and hydrophobic forces [27, 28, 31, 33–36, 38–46]. These interactions may be mediated by intrinsically disordered regions (IDRs) or low complexity motifs that become ordered when bound to RNA [40, 44, 45, 47] or other biomolecules. Cytoplasmic examples of BMCs include stress granules and P bodies [48, 49], and nucleoli, Cajal bodies, and transcriptional condensates have been described as nuclear BMCs [50]. Nuclear processes that are driven by concentrating critical molecules in BMCs include transcription, splicing, DNA repair, and chromatin modification [44, 50–52]. If phase transition states become dysregulated and progress from liquid to solid, pathological gels and fibrils can form, resulting in disease states [31, 53, 54].

Numerous viruses have been reported to use phase separation as a mechanism to facilitate virus replication. For example, measles, rabies, influenza, vesicular stomatitis virus, and adenovirus viral ribonucleoprotein (vRNP) replication complexes utilize LLPS as a mechanism to concentrate viral components within cells [55–61]. Among retroviruses, human immunodeficiency virus type 1 (HIV-1) Gag has been shown to form BMCs, with the NC protein appearing to be a major driver of condensate formation [62]. Based on computational predictions indicating that RSV Gag contains IDRs, and in light of the observation that RSV Gag forms foci in different subcellular compartments, we tested the hypothesis that RSV Gag possess biophysical properties of liquid-like, phase-separated BMCs. Our experimental results suggest that Gag is finely tuned to form BMCs with differing characteristics during the assembly pathway from the nucleus to the plasma membrane. Further evidence for the importance of phase transitions in RSV biology stem from our observations that the biophysical properties of RSV Gag are sensitive to concentration and to mutations that lead to alterations in the liquid-like properties of the protein.

## Results

### RSV Gag contains IDRs and forms phase-contrasted nuclear foci

We previously observed that RSV Gag forms spherical foci in the nucleus, cytoplasm, and at the plasma membrane, representing each of the subcellular compartments involved the assembly pathway [3, 22]. Approximately 20% of wild-type RSV Gag localizes to the nucleus under steady-state conditions [17, 20], and interfering with CRM1-mediated nuclear export by mutating the p10 NES (e.g., mutant Gag.L219A), treating cells with leptomycin B (LMB), or overexpressing dominant-negative mutants of nucleoporins Nup98 or Nup214, causes nuclear Gag foci to accumulate in size and number [17, 19]. The spherical appearance of Gag foci, as well as the observation that they are enhanced with increased concentration, suggested that these protein-rich foci could form BMCs, as described for other viral and cellular complexes. Others have noted that RSV Gag contains unstructured domains [63], and the presence of IDRs has been described in the HIV-1 Gag protein [62]. To examine more specifically whether the RSV Gag polyprotein contains canonical IDRs, we used PONDR (Predictor of Natural Disordered Regions; www.pondr.com) to analyze the amino acid sequence (Fig. 1A). We identified two prominent IDRs: one encompassing the C-terminal portion of MA plus the adjacent p2 and p10 domains; and the other extending throughout the NC domain.

To determine whether nuclear foci formed by wild-type or mutant Gag proteins form phase-contrasted complexes--as described for other IDR-containing proteins (reviewed in [64])—we performed confocal fluorescence microscopy and differential interference contrast (DIC) imaging (Fig. 1B-E). The wild-type Gag protein forms foci in the cytoplasm, at the plasma membrane, and in the nucleus (outlined in white) as previously reported (Fig. 1B, Gag-SNAPTag fusion protein, green), with nuclear foci demonstrating phase contrast overlapping with the fluorescent signal of the protein (Fig. 1B; arrows). Treatment of cells with LMB or co-expression of a dominant-negative Nup214 protein (NP214-DsRed, red) resulted in accumulation of numerous nuclear Gag-SNAPTag foci that exhibited phase contrast (Fig. 1C and D, respectively) [19]. The p10 NES mutant Gag.L219A-CFP also formed prominent nuclear complexes that corresponded to phase-contrasted foci in the DIC images. As noted previously, Gag nuclear foci appear larger and more numerous when nucleocytoplasmic transport is inhibited, either by expression of a dominant-negative NP214, treatment with LMB, or mutation of the p10 NES, indicating that the formation of Gag nuclear foci is concentration-dependent, a characteristic of BMCs [19]. Together, these data suggest that Gag meets two criteria of BMCs: the presence of IDRs and the ability to form spherical foci that are visible under DIC imaging.

### Recombinant Gag proteins form phase-contrasted droplets *in vitro*

Another characteristic of proteins that form BMCs is their ability to form liquid-like droplets *in vitro* that demonstrate separation from the surrounding milieu [27, 34–36]. To determine whether RSV Gag proteins were capable of forming phase-separated droplets in vitro, recombinant full-length Gag.ΔPR (WT Gag), Gag.L219A, and Gag deletion mutants were highly purified from *E. coli* in the absence of nucleic acids, labeled with Alexa 488, and mixed with a crowding agent to a final protein concentration of 5 µM, 10 µM, or 20 µM (Fig. 2). Under all conditions, full-length Gag.ΔPR (WT, Fig. 2A) formed droplets that demonstrated phase-contrast by DIC. Based on the prediction that IDRs were located in the MAp2p10 and NC domains of Gag (Figure 1A), we examined various truncation mutants to determine whether they possessed the ability to form *in vitro* droplets. Although each construct was capable of forming droplets in vitro, the number and size of droplets varied considerably among the purified proteins.

**Figure 2:**
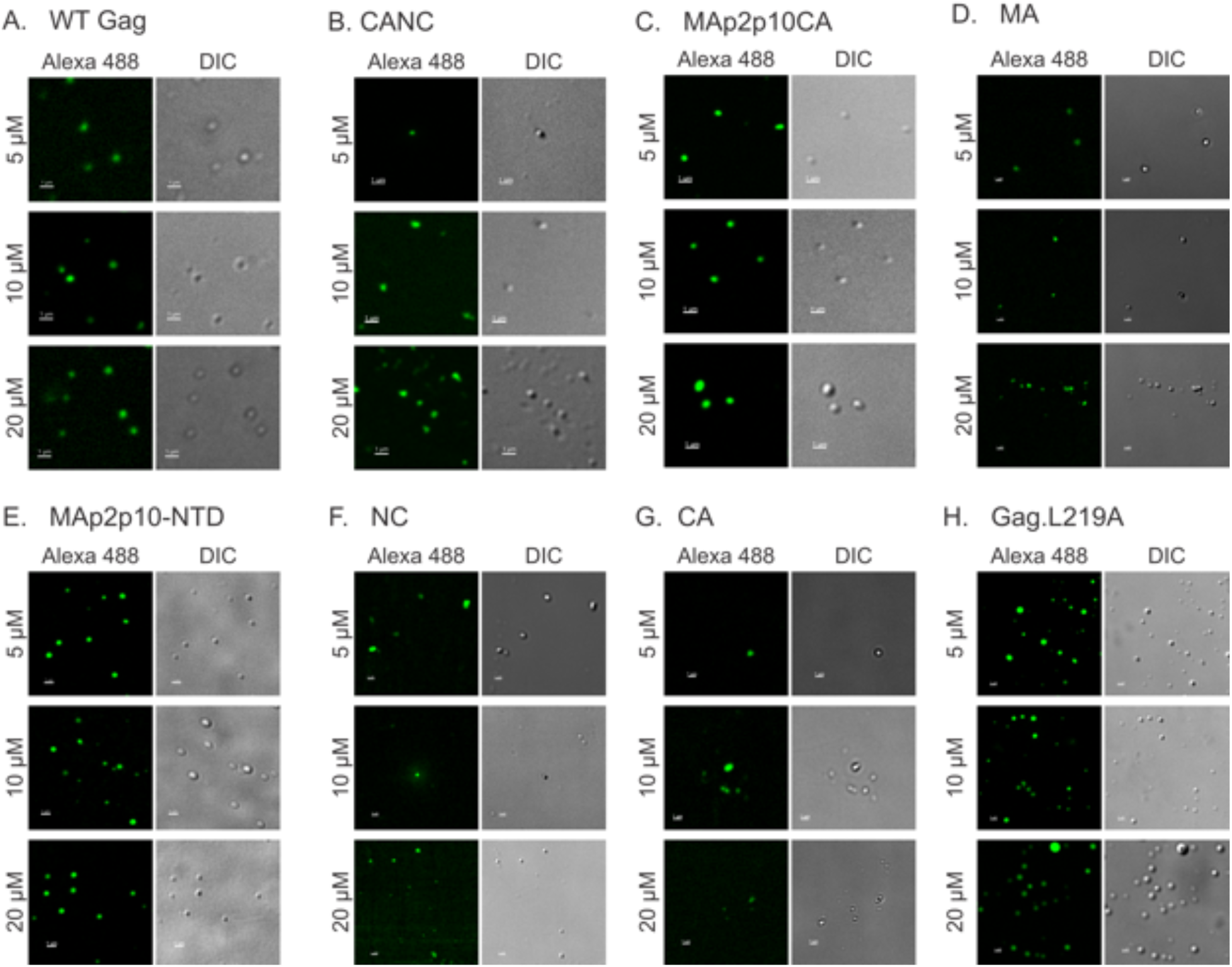
Full-length and mutant purified RSV Gag proteins form *in vitro* droplets. Purified proteins were labeled with Alexa Fluor 488 and imaged via confocal microscopy at 7x zoom. Protein concentration is indicated to the left. All proteins formed droplets under these conditions, to varying degrees. Scale bar= 1 µm. DIC was relied upon for visualization and analysis of NC and CA complexes due to inefficient labeling with Alexa Fluor dye (see Methods).

To quantitate the differences in size and number at each concentration, low magnification (1x) fluorescence images of each construct were analyzed using the spot function in the Imaris imaging analysis program (Fig. 2 and 3; Supplemental Tables 1 and 2). For NC and CA, which lack abundant primary amines for labeling (see Methods) making their fluorescence difficult to visualize, DIC images were used rather than fluorescence images. Only spherical droplets were counted to avoid skewing the measured diameter of the droplets. WT Gag formed droplets that exhibited a concentration-dependent increase in size (p<0.0001 for each concentration), with 10 µM yielding the highest number of droplets (p<0.001 compared to 5 µM and p=0.0047 compared to 20 µM). The observed decrease in droplet number at a concentration of 20 µM could be due to nucleation of smaller droplets into less abundant, larger droplets.

**Figure 3:**
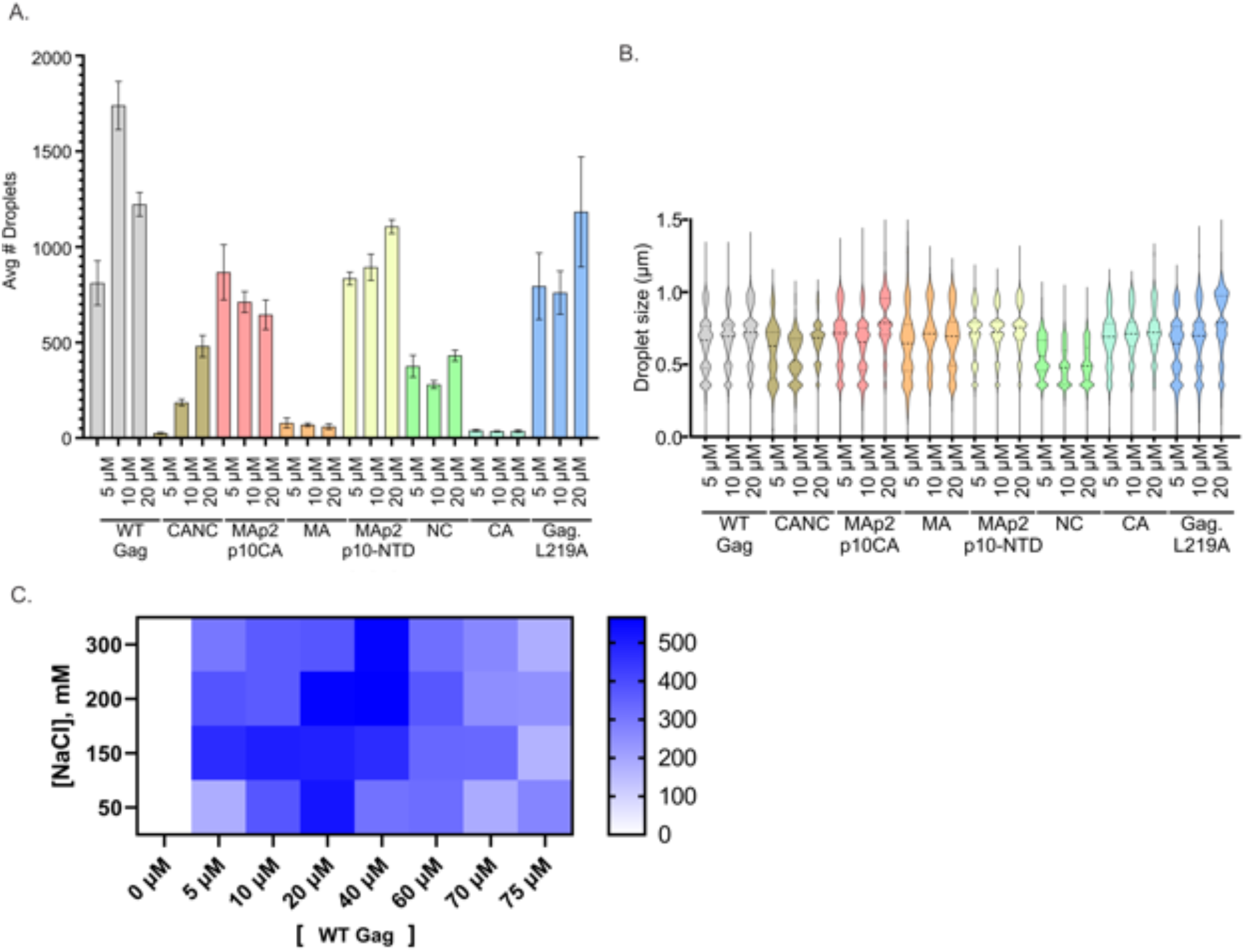
Analysis of *in vitro* droplet size, number, and protein/salt concentrations required for droplet formation. (A) and (B) 10 low magnification fields were captured for each protein and concentration. The Imaris spot function was used to count the number of droplets and estimate the diameter of each droplet. Non-spherical droplets and those at the edge of the field were excluded so as not to skew the diameter measurement. Fluorescent images were subjected to Gaussian filtering to remove background fluorescence. The fluorescence channel of all proteins was utilized for analysis with the exception of NC and CA, due to inefficient fluorescent labeling (see Methods). For most proteins, the average number of droplets per field varied for each concentration (A). A violin plot was used to display the distribution of droplet sizes for each condition (B). To determine the optimal conditions for *in vitro* droplet formation for WT Gag, a phase diagram was generated by comparing various protein and NaCl concentrations and counting the number of droplets present under each condition (C). The protein was unlabeled, and droplets visualized by DIC were counted using the Imaris spot function. White indicated zero droplets, with increasingly darker shades of blue representing the indicated number of droplets averaged from ≥5 fields per condition. Based upon this analysis, NaCl concentration of 150 nM and Gag concentration of 10 µM to 20 µM appeared to provide the optimal conditions for *in vitro* droplet formation.

The largest predicted IDR in Gag was in MAp2p10, so we deleted this region to determine whether it was necessary for the formation of *in vitro* droplets (CANC, Figure 2B). Compared to WT Gag, CANC formed significantly fewer and smaller spherical droplets (number vs WT: p<0.0001 for all concentrations; size vs WT: 5 µM p= 0.0009, 10 µM and 20 µM p<0.0001), suggesting that MAp2p10 contributes to the efficiency of *in vitro* droplet formation. The other IDR is predicted to be located in the NC domain (Figure 1A), so we next deleted this region from Gag and assessed droplet number and size. Deletion of NC led to the formation of droplets that were larger at 5 µM and 20 µM concentrations, yet smaller for 10 µM compared to WT Gag at the same concentrations (p<0.0001 for all concentrations).

To further determine the sufficiency of individual Gag domains to form droplets, we performed the same *in vitro* droplet assay using segments of Gag. We first examined whether MA alone was capable of forming droplets, as it contains only a small portion of the IDR (Figure 2D). The size of MA droplets was similar at 5 µM, larger at 10 µM, but smaller at 20 µM compared to WT Gag at the same concentrations (p<0.0001 at 10 and 20 µM concentrations). However, the number of droplets formed by MA was greatly decreased compared to WT Gag (<100 droplets per field) and did not vary with differing protein concentration, suggesting that MA alone does not efficiently undergo LLPS to form droplets; alternatively, the observed complexes could be formed by other types of interactions. Although we wished to examine the propensity of MAp2p10 to form droplets, it was insoluble when extracted from *E. coli* (data not shown), possibly due to its disorder. However, by adding the structured N-terminal domain of CA (NTD), the protein was well-behaved. MAp2p10-NTD efficiently formed droplets that increased in size with increasing protein concentration and were significantly larger than WT at the corresponding protein concentrations (p<0.0001 at all concentrations). Although MAp2p10- NTD formed fewer droplets than WT Gag, the difference was only significant at 10 µM (p<0.0001), again suggesting that MAp2p10 plays a major role in Gag’s ability to undergo phase separation. The slightly decreased droplet count of MAp2p10-NTD compared to WT Gag suggests that the NC domain IDR also contributes to Gag droplet formation in the context of full length Gag. Although NC alone produced fewer and smaller droplets at each protein concentration compared to WT Gag, it made more droplets compared to MA alone, suggesting that NC is sufficient to form BMCs and likely contributes to Gag droplet formation *in vitro*.

CA does not contain an IDR, but contains structured N- and C-terminal domains and plays an important role in the formation of the viral capsid in mature virions [63]. Therefore, we set out to determine whether it is capable of forming phase-separated droplets in our *in vitro* assay. Under these conditions, CA produced droplets very inefficiently, with the lowest number of complexes formed of all constructs tested. Interestingly, the number of CA complexes did not increase with increasing protein concentration. These results suggest that CA does not form BMCs, but instead is consistent with previous findings indicating that CA forms a stable ordered complex required for capsid structure in the mature virion [65].

Based on our observation in cells that mutating the Gag p10 NES (Gag.L219A) results in large nuclear foci suggestive of BMCs [19], we asked whether Gag.L219A forms droplets *in vitro* (Figure 2H). To our surprise, droplet size for this protein increased significantly as protein concentration increased (p<0.0001 at all concentrations). However, the number of droplets was not significantly different among each protein concentration and was similar to WT Gag at all concentrations except 10 µM. At this concentration, Gag.L219A formed fewer droplets compared to WT Gag (p<0.0001), suggesting that this mutation in the p10 sequence within the IDR does have a minor effect on *in vitro* droplet formation.

To define the phase diagram for WT Gag, the protein and salt (NaCl) concentrations were varied (Figure 3C). Assays were conducted in the range of protein concentrations from 5-75 µM and NaCl concentrations varying from 50-300 mM using unlabeled purified WT Gag protein and imaged using DIC. Droplets were imaged at low magnification and the average number of droplets in ≥5 fields were counted using the Imaris spot function. In the heat map, white indicates zero droplets with increasing intensity of blue color showing more droplets with a maximum of dark blue, which contained over 500 droplets. As expected for a protein undergoing phase separation, Gag droplet formation was sensitive to protein and salt concentration. Gag did not form droplets efficiently at the lowest concentrations of protein and NaCl (5 µM, 50 mM, respectively); droplet formation peaked at mid-range concentrations of 10-40 µM and 150 mM NaCl; and fewer particles formed at the highest protein concentrations tested (>40 µM). For WT Gag, 150 mM appeared to be the ideal salt concentration for most protein concentrations, with 10-20 µM being the optimal protein concentration.

### Gag droplets undergo fusion and fission, demonstrating liquid-like properties

To determine whether Gag droplets formed *in vitro* were dynamic (Figure 4), 20 µM of unlabeled WT Gag or Gag.L219A protein was used in an *in vitro* droplet assay and imaged directly after mixing and for every 10 seconds for a defined period under brightfield for WT Gag (Figure 4A; Supplemental Movie 1, black circle) or DIC for the p10 NES mutant Gag.L219A (Figure 4B; Supplemental Movie 2, black circles). For both proteins, fusion between two droplets (white arrows) occurred over time.

**Figure 4:**
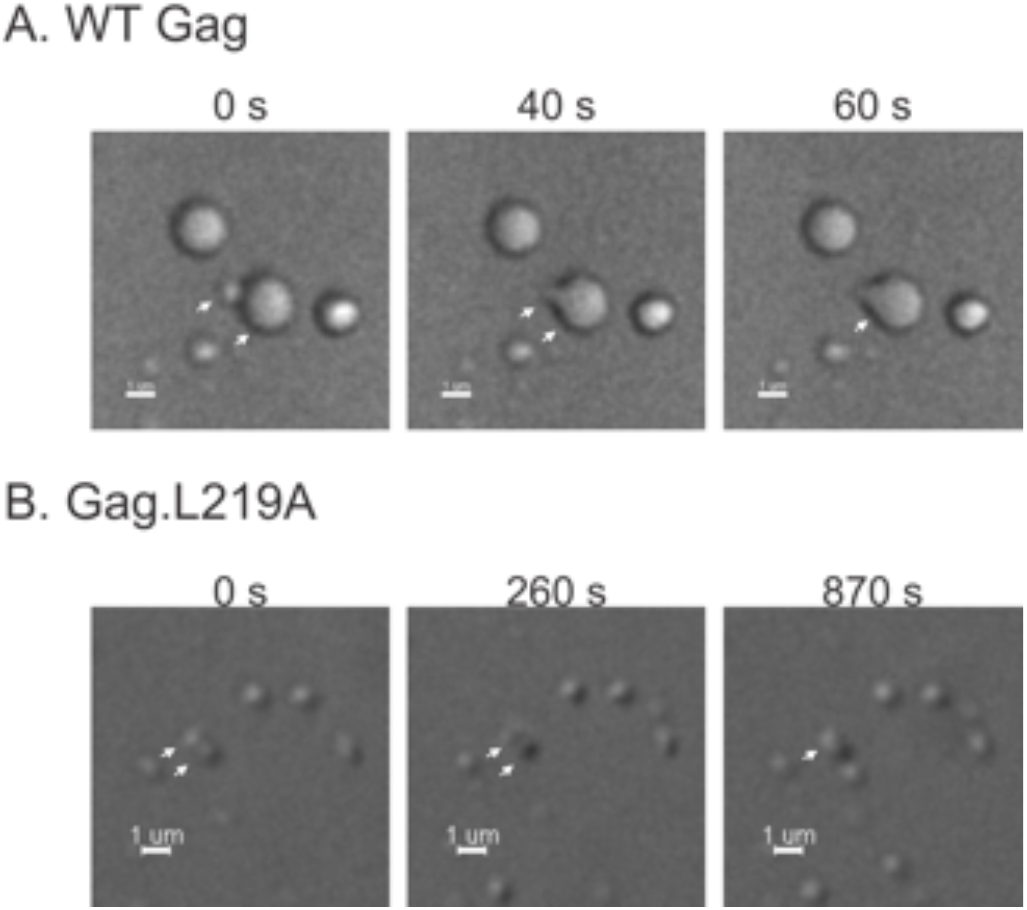
*In vitro* droplet fusion. 20 µM of unlabeled protein complexes composed of either WT Gag or Gag.L219A were imaged immediately after mixing. Complexes were imaged every 10 seconds under either bright field (WT) or DIC (Gag.L219A). (A) A small Gag droplet was seen fusing with a larger droplet (designated by white arrows) over time. (B) Two small Gag.L219A droplets (white arrows) fused into a single droplet over time. Scale bar= 1 µm.

To examine fusion and fission *in vivo*, living QT6 cells expressing Gag-SNAPTag were imaged at 1 frame/minute (Figure 5A, plus Supplemental Movies 3a, 3b, and c). For better visualization of the foci in each cellular compartment, a separate movie was generated with the histogram differently adjusted for the nucleus (Supplemental Movie 3a), cytoplasm (Supplemental Movie 3b), and plasma membrane (Supplemental Movie 3c). In the first frame shown in Fig. 5A (t = 2 min), two individual separate nuclear foci are indicated by white arrows (nucleus outlined in dashed white line; see also zoomed in image of boxed region below and Supplemental Movie 3a, puncta enclosed in white circle). At the next time point (t = 3 min) the individual nuclear puncta fused into a single focus. Throughout the still images and supplemental movies of this cell, several foci were observed undergoing fusion and fission in the cytoplasm (yellow arrows shown in main image and in boxed area zoomed in below; yellow circles in Supplemental Movie 3b) and at the plasma membrane (magenta arrows in main image and in boxed area zoomed in below; magenta circle in Supplemental Movie 3c), indicating that Gag foci are dynamic in all major subcellular compartments. To examine the dynamic properties of the nuclear-localized Gag.L219A-YFP NES mutant, time-lapse live cell imaging was performed under similar conditions (Figure 5B; Supplemental Movie 4, black circles). We observed several fusion events between nuclear foci which were separate at the beginning of the imaging sequence and subsequently fused into single foci after 2 min (white arrows). Taken together, these data demonstrate that WT Gag and Gag.L219A exhibit liquid-like properties, evidenced by the formation of spherical droplets that undergo fusion and fission *in vitro* and *in vivo*.

**Figure 5:**
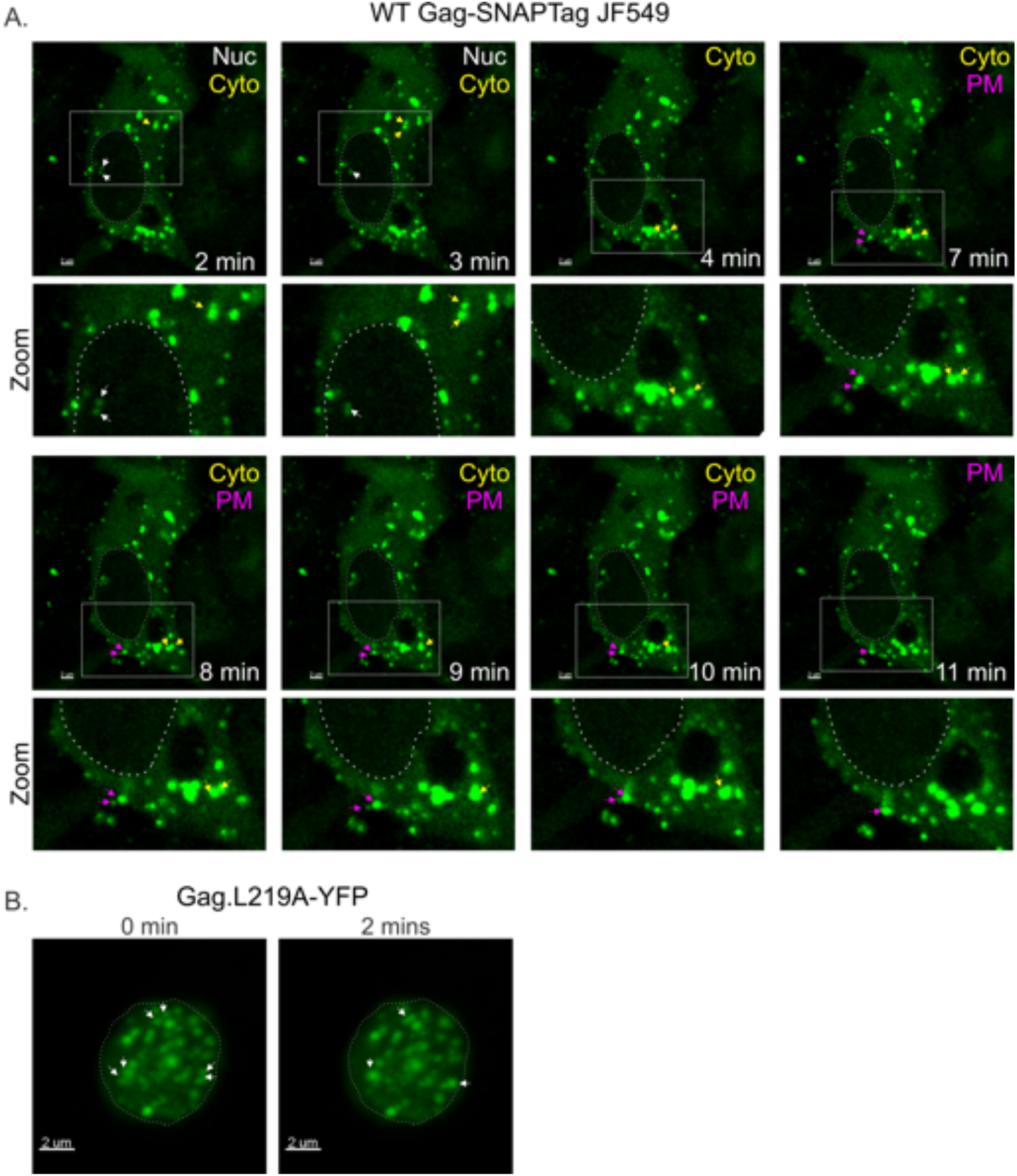
*In vivo* droplet fusion and fission of WT Gag and Gag.L219A. One characteristic of BMCs is that they are dynamic and can undergo fusion or fission over time. (A) Using a single confocal z-plane, live-cell timelapse imaging was conducted (1 frame/minute) in a QT6 cell expressing WT Gag-SNAP-tag bound to SNAP-ligand conjugated to JF549. The nucleus was outlined in a white dashed line. Several WT Gag-SNAP-tag foci were observed to undergo fusion or fission in different cellular compartments: nucleus (foci indicated by white arrows), cytoplasm (yellow arrows), and at the plasma membrane (magenta arrows). Droplets of interest were also displayed at higher magnification (inset). See also Supplemental Movies 3a, 3b, and 3c for timelapse images. (B) A single optical Z-plane of a QT6 cell expressing the Gag.L219A- YFP nuclear export mutant was imaged using confocal timelapse microscopy at one frame every minute. Over a period of two minutes, three different pairs of droplets underwent fusion. The nucleus is outlined with a white dotted-line. Scale bar= 2 µm. See also Supplemental Movie 4 for timelapse images.

### Measuring protein dynamics using fluorescence recovery after photobleaching (FRAP)

To determine whether Gag proteins located in spherical foci exchange freely with molecules in the surrounding environment, we performed FRAP experiments of Gag-YFP foci in the nucleus, cytoplasm, and along the plasma membrane (Fig. 6), as previously described [22]. For these experiments, the entire focus was bleached irreversibly and fluorescence was monitored to determine the half-time of recovery (t½) and the mobile fraction. Foci from numerous cells (range 14-23) were sampled and the means for each value were calculated (Figure 7D and E, Tables 1-4). For WT Gag foci in the nucleus, the half-time of recovery was rapid (1.25 ± 0.22 seconds) and the mobile fraction was 25 ± 2 %. Gag foci in the cytoplasm had a slightly longer t½ of recovery (1.33 ± 0.13 seconds) and a significantly higher mobile fraction (33 ± 2.9%, *p=0.0473). For Gag foci located at the plasma membrane, the t½ was very similar to the cytoplasmic foci at 1.34 ± 0.14 seconds, and the mobile fraction 26 ± 2.9%. These data suggest that Gag foci exchange rapidly—but only to a moderate degree with surrounding molecules in all three locations—with nuclear and plasma-membrane localized Gag molecules appearing to be more fixed in the complex, potentially due to their interaction with a particular set of host factors involved in their transport or localization.

**Figure 6:**
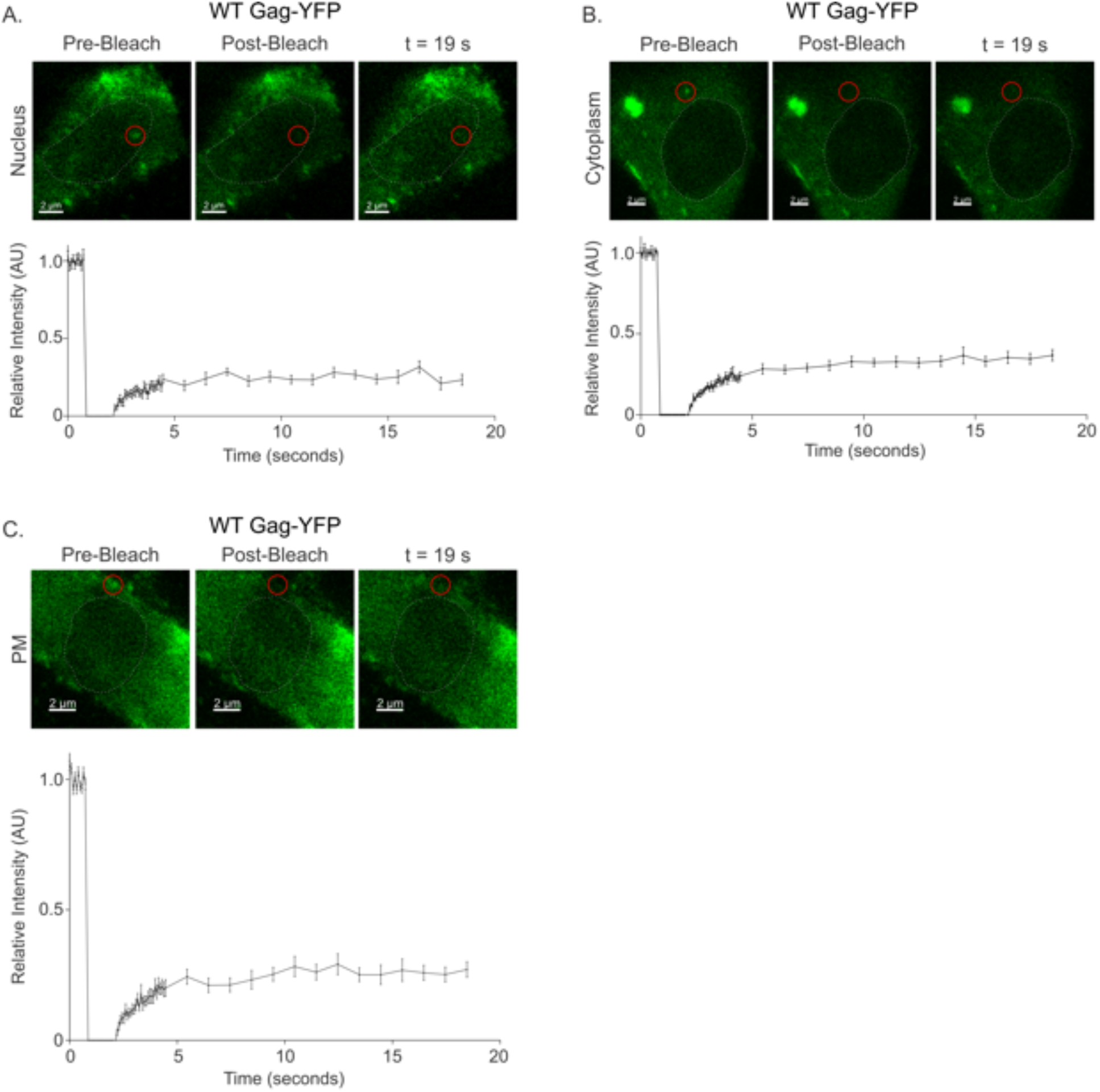
FRAP analysis of WT Gag foci in the nucleus, cytoplasm, and at the plasma membrane. WT RU5.Gag-YFP foci (red circles) were photobleached and the half-time (t1/2) and fraction of recovery was measured for approximately 19 seconds. (A) Nuclear foci (N=14), (B) cytoplasmic foci (N=18), (C) plasma membrane foci (N=17) were analyzed. The graphs show the average recovery graph with error bars for the indicated number of foci. Scale bar= 2 µm. The nucleus was outlined in a white dashed line. See also Tables 1, 3, and 4 for quantification and statistical analysis.

**Figure 7:**
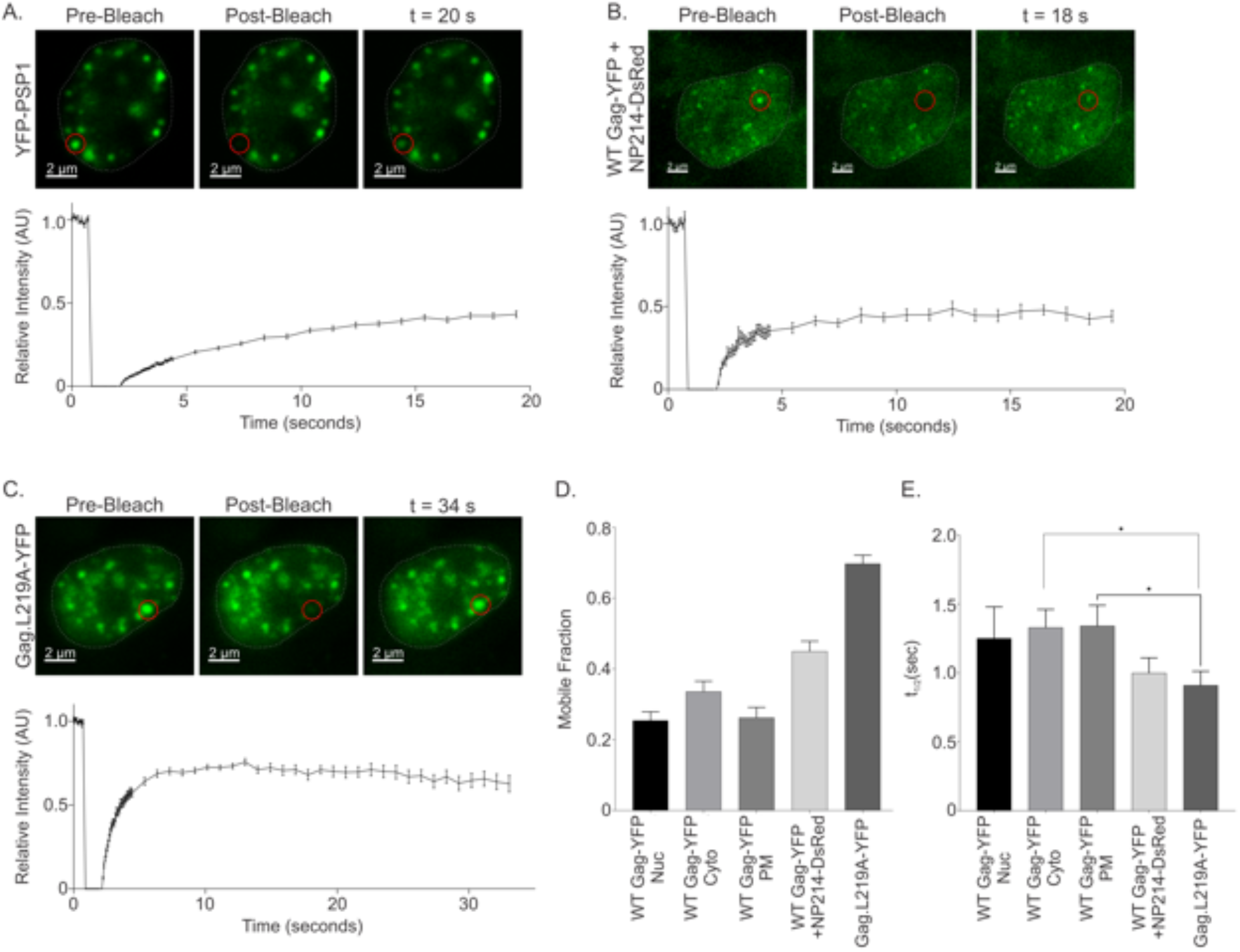
FRAP analysis of PSP1 and nuclear restricted Gag foci. (A) YFP-PSP1 (N=23), a paraspeckle protein, served as a control for the FRAP experiments. (B) To block WT Gag nuclear export and increase the nuclear concentration of Gag protein, a dominant negative nucleoporin Nup214 (NP214-DsRed) was co-expressed with WT RU5.Gag-YFP (N=15). (C) The nuclear export mutant RU5.Gag.L219A-YFP (N=20) formed bright nuclear complexes that recovered quickly. Scale bar= 2 µm. (D) and (E) Graphs presenting the average mobile fraction and t1/2 (sec) comparisons for each Gag condition. The nuclear Gag foci formed with NP214 co-expression or Gag.L219A foci recovered with a significantly shorter t1/2 and higher mobile fraction compared to WT Gag foci shown in Figure 6. See also Tables 2, 3, and 4 for quantification and statistical analysis.

**Table 1:**
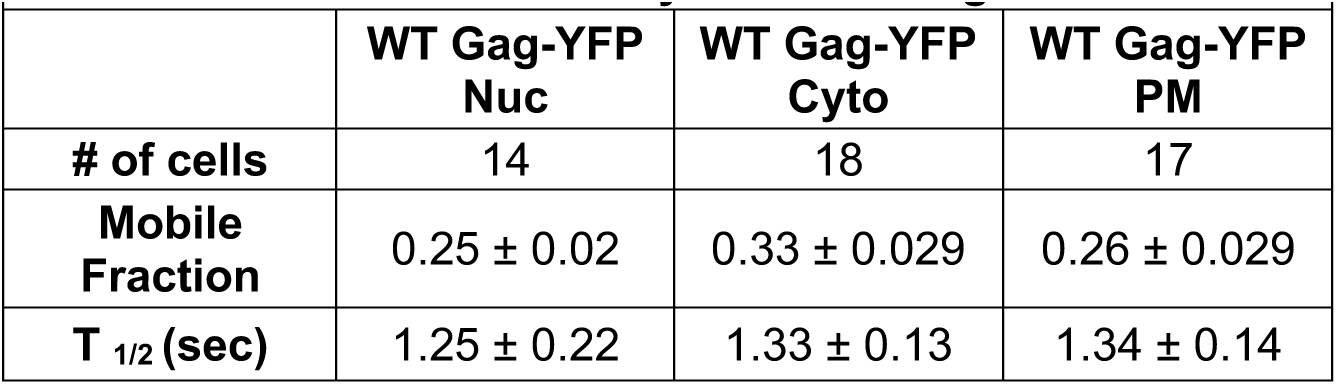
FRAP mobile fraction and t1/2 (sec) analysis of WT Gag in the nucleus, cytoplasm, and at the plasma membrane.

**Table 2:**
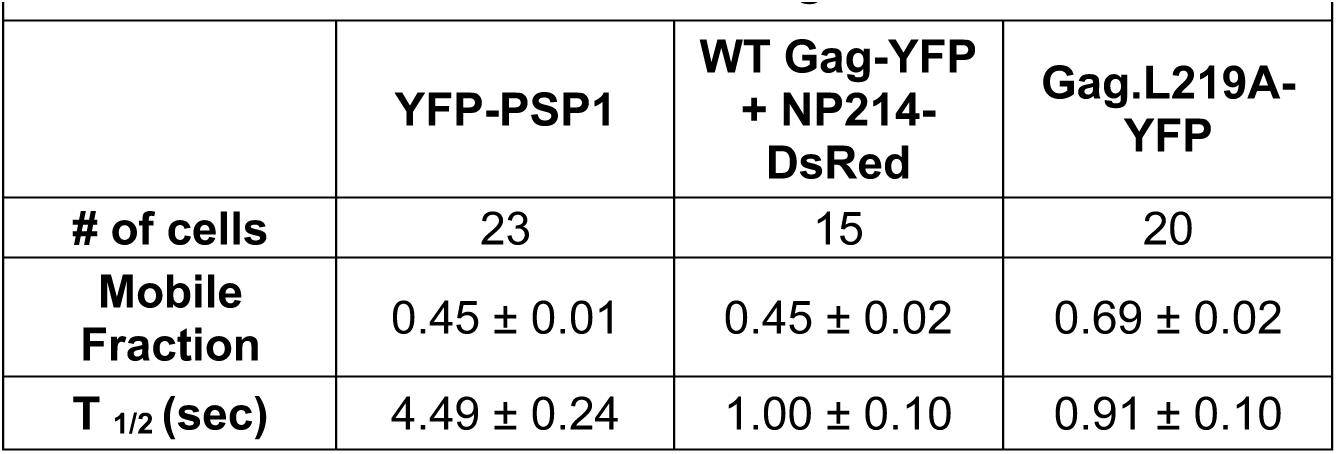
FRAP mobile fraction and t1/2 analysis of YFP-PSP1 and nuclear restricted Gag.

**Table 3:** FRAP mobile fraction comparison of WT and mutant Gag proteins.

| **Table 3: Mobile Fraction Comparison** |  |  |  |  |  |
| --- | --- | --- | --- | --- | --- |
|  | **Means** | **WT Gag-YFP Nuc** | **WT Gag-YFP Cyto** | **WT Gag-YFP PM** | **WT Gag-YFP + NP214-DsRed** |
| **WT Gag-YFP Nuc** | 0.25 ± 0.02 |  |  |  |  |
| **WT Gag-YFP Cyto** | 0.33 ± 0.029 | *<br>0.0473 |  |  |  |
| **WT Gag-YFP PM** | 0.26 ± 0.029 |  |  |  |  |
| **WT Gag-YFP + NP214-DsRed** | 0.45 ± 0.02 | ****<br><0.0001 | **<br>0.0093 | ****<br><0.0001 |  |
| **Gag.L219A-YFP** | 0.69 ± 0.02 | ****<br><0.0001 | ****<br><0.0001 | ****<br><0.0001 | ****<br><0.0001 |

Unless indicated by \*, the comparisons were not statistically significant

**Table 4:** FRAP t1/2 (sec) comparison of WT and mutant Gag proteins.

Unless indicated by \*, the comparisons were not statistically significant
| **Table 4: $t_{1/2}$ Comparison** |  |  |  |  |  |
| --- | --- | --- | --- | --- | --- |
|  | **Means** | **WT Gag-YFP Nuc** | **WT Gag-YFP Cyto** | **WT Gag-YFP PM** | **WT Gag-YFP + NP214-DsRed** |
| **WT Gag-YFP Nuc** | 1.25 ± 0.22<br>secs |  |  |  |  |
| **WT Gag-YFP Cyto** | 1.33 ± 0.13<br>secs |  |  |  |  |
| **WT Gag-YFP PM** | 1.34 ± 0.14<br>secs |  |  |  |  |
| **WT Gag-YFP + NP214-DsRed** | 1.00 ± 0.10<br>secs |  |  |  |  |
| **Gag.L219A-YFP** | 0.91 ± 0.10<br>secs |  | *<br>0.0149 | *<br>0.0190 |  |

1180 Unless indicated by \*, the comparisons were not statistically significant

We next assessed the effect of inhibiting WT Gag nuclear export on its exchangeability by co-expressing the trans-dominant NP214 mutant (Figure 7B), which resulted in an increase in the number and size of Gag nuclear foci. In these FRAP experiments, the t½ = 1.0 ± 0.10, indicating that the protein was more dynamic, and the mobile fraction was higher at 45 ± 2% (compared with: WT Gag-YFP Nuc, ****p<0.0001; WT Gag-YFP Cyto. ** p=0.0093; WT Gag-YFP PM, ****p<0.0001). By comparison, the Gag nuclear export mutant (Gag.L219A; Figure 7C) had the largest nuclear foci, the highest mobile fraction (69 ± 2%, ****p<0.0001 compared to all conditions), and the most rapid exchange kinetics with t½ = 0.91 ± 0.10 seconds (Compared to WT Gag-YFP Cyto, *p=0.0149; WT Gag-YFP PM, *p=0.0190). In both cases, increasing the intranuclear concentration of Gag resulted in more dynamic behavior of the protein, suggesting it adopted more liquid-like behavior in a concentration-dependent manner. As a control, we expressed PSP1 protein (Figure 7A), which is known to form BMCs as a component of paraspeckles [66]. This protein had a t½ of 4.49 ± 0.24 seconds and mobile fraction of 45 ± 1% in QT6 cells. This t½ for PSP1 was in the same range as that of a previously reported value (6.42 seconds), although the mobile fraction we measured was decreased [67], likely due to differences in cell type and in endogenous vs transient expression.

### RSV Gag IDRs contribute to condensate formation in optoDroplets

To determine which regions of the RSV Gag protein contribute to condensate formation using an *in vivo* IDR assay in HEK293T cells, we generated constructs consisting of mCherry-tagged proteins derived from RSV Gag (RSV Gag ΔPR, MAp2p10, CA, and NC) fused to the photoactivatable domain of the CRY2 protein from *Arabidopsis thaliana* [68–71]. The optoDroplet system leverages the blue light-activated clustering of CRY2 to assess the propensity of protein sequences to form BMCs, as evidenced by real-time droplet formation observed via live cell confocal microscopy (68-71).

As a baseline, CRY2oligo-mCherry alone had minimal clustering when activated by blue light (Figure 8A, Supplemental Movie 5). As reported previously, the FUS.IDR caused marked clustering of CRY2oligo when stimulated using blue light (Figure 8B, Supplemental Movie 6) [69]. Of the RSV Gag proteins linked to CRY2oligo-mCherry, RSV Gag ΔPR, MAp2p10, and NC all exhibited clustering with foci formation upon blue light exposure (Figures 8C-E, Supplemental Movies 7-9). In the absence of activation, RSV Gag ΔPR.CRY2oligo-mCherry and MAp2p10.CRY2oligo-mCherry appeared to be diffuse, but upon blue light exposure, these two proteins formed discrete foci at the perimeter of the cells, associated with the plasma membrane (Figures 8C and D, Supplemental Movies 7 and 8). This localization pattern was consistent with the plasma membrane-binding and targeting properties of the Gag MA domain [72]. The NC.CRY2oligo-mCherry protein was diffuse in the nucleus in the absence of activation, and once stimulated with blue light, it formed numerous, discrete foci in the nucleoplasm (Figure 8E, Supplemental Movie 9). In contrast, CA, which does not contain a predicted IDR, failed to induce condensation of CRY2oligo (Figure 8F, Supplemental Movie 10). These results suggest that the IDRs in both MAp2p10 and NC are sufficient to drive to BMC formation in a cell-based assay.

**Figure 8:**
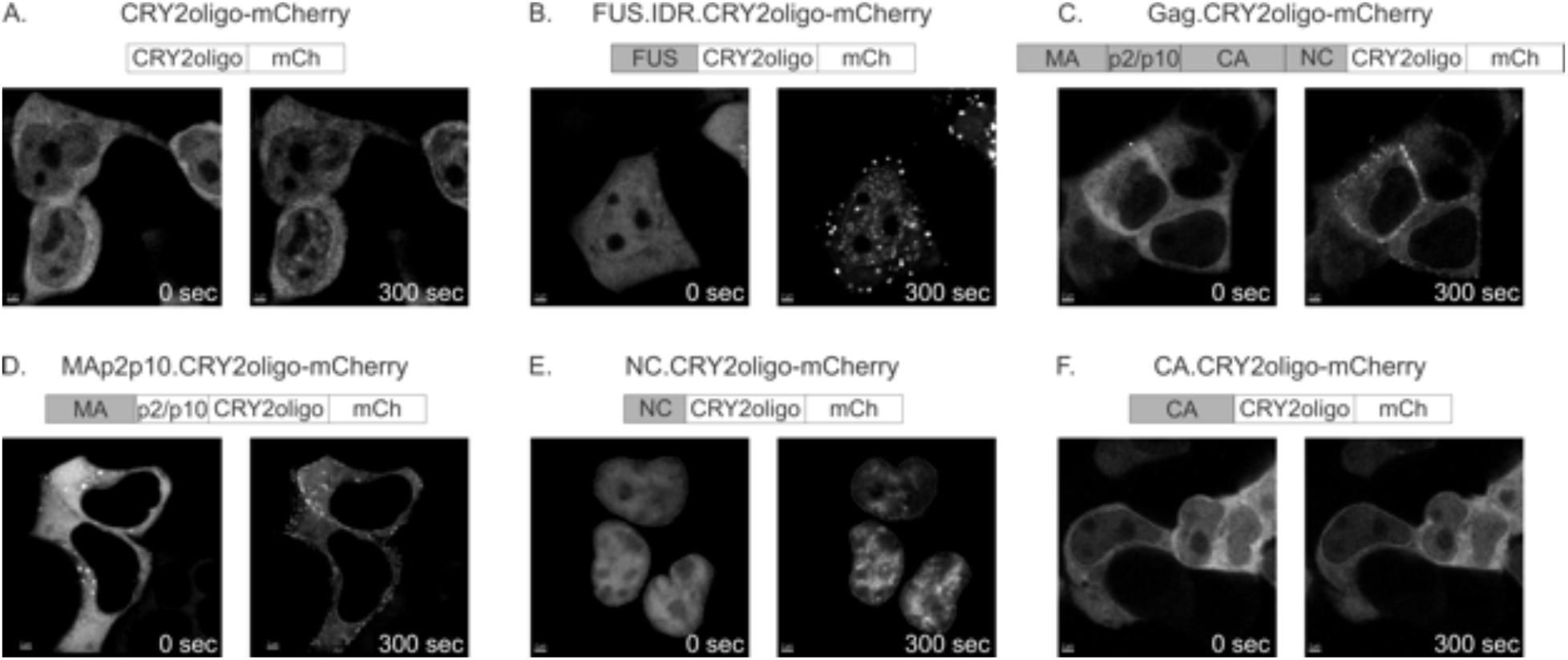
OptoDroplet Assay of Gag IDRs. To determine whether either WT Gag or individual domains of Gag could form condensates in living cells, they were fused to CRY2oligo, which induces droplet formation when illuminated with blue light, and mCherry to allow for fusion protein visualization. Living HEK293T cells expressing the indicated constructs were imaged every 5 seconds for 300 seconds. Blue light illumination occurred 30 seconds into the time course. A) CRY2oligo-mCherry formed minimal clusters with blue light illumination. B) FUS.IDR.CRY2oligo-mCherry had strong clustering with blue light. C) Gag.CRY2oligo-mCherry and D) MAp2p10.CRY2oligo-mCherry formed plasma membrane localized condensates. E) NC.CRY2oligo-mCherry formed numerous nuclear droplets. F) CA.CRY2oligo-mCherry, which does not contain an IDR, did not cluster with blue light illumination. Scale bar= 2 µm. Nuclei were outline by white dashed lines. See also Supplemental Movies 5, 6, 7, 8, 9, and 10.

### Exchangeability of canonical IDRs from FUS and HNRNPA1 with RSV Gag IDRs restored condensate formation in cells

The data presented thus far suggest that RSV Gag undergoes LLPS to form BMCs that are primarily driven by IDRs in MAp2p10 and NC. To test whether classically defined IDRs from two well-studied proteins, FUS and HNRNPA1, could replace the IDRs in Gag, we produced chimeric proteins for a set of gain-of-function experiments in QT6 cells (Figure 9). In the absence of the NC domain, Gag was distributed throughout the cytoplasm due to the loss of the nuclear localization signal [16, 17], and the deletion mutant fails to form foci as reported previously [17], including at the plasma membrane (compare Figures 1B and 9A). Substitution of the FUS IDR for the NC sequence in Gag restored formation of foci in the nucleus, cytoplasm, and at the plasma membrane (Figure 9B), demonstrating that the presence of an IDR in the NC region is sufficient to recreate the typical pattern of Gag distribution.

**Figure 9:**
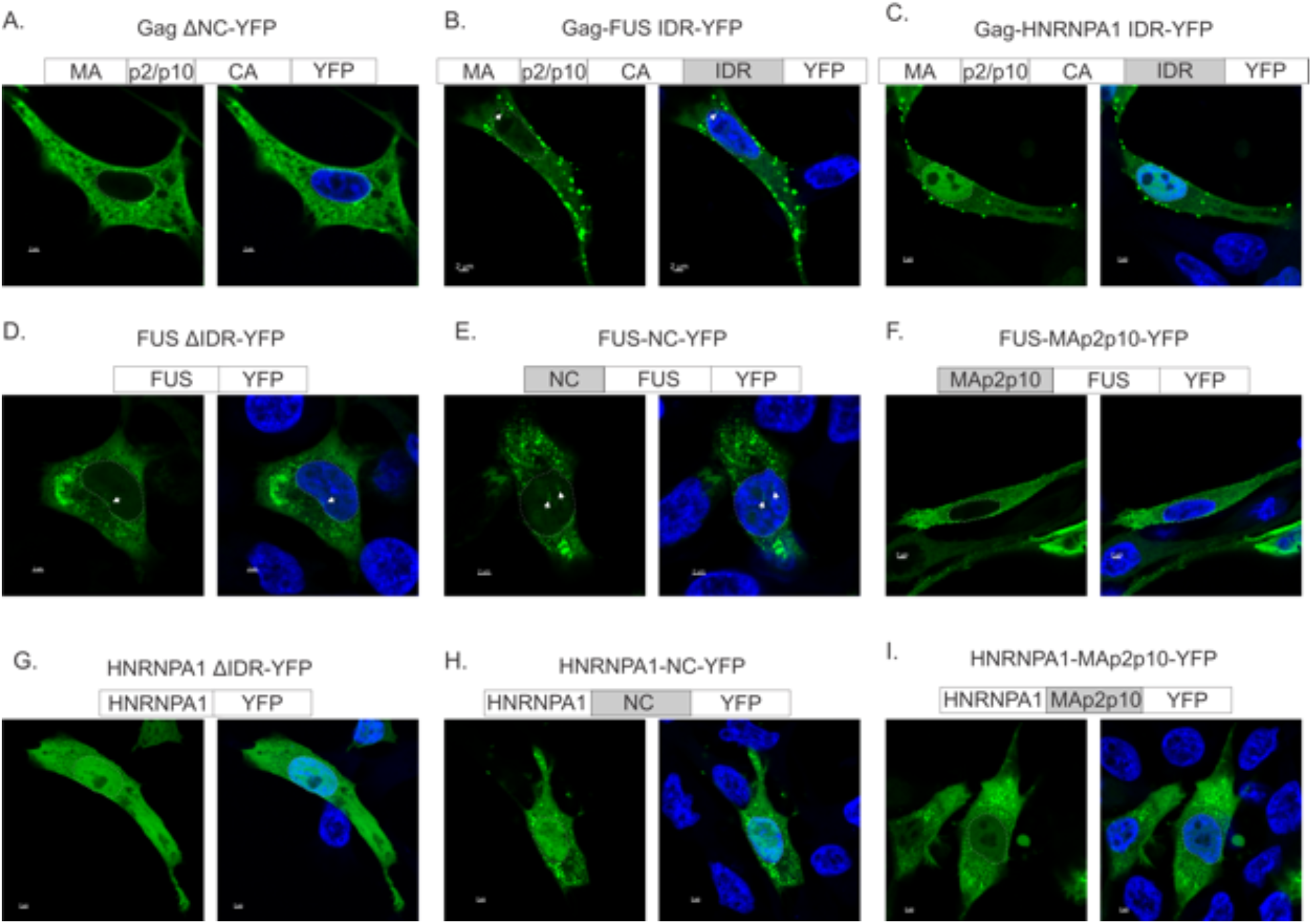
Chimeras of Gag and cellular protein IDRs. To determine whether the putative IDRs from either Gag or cellular proteins with well-characterized IDRs (FUS and HNRNPA1) could rescue the ability of mutants lacking IDRs to form droplets, chimeras were created. Schematic diagrams of the chimeras were placed above the microscopic images with IDRs shaded gray. Representative confocal microscopy images were shown for each construct. (A) In the absence of the NC domain (Gag ΔNC-YFP), Gag remained diffuse in the cytoplasm and lost the ability to form foci. (B) Replacing the NC sequence with the FUS IDR allowed Gag to form foci once again (RU5.Gag-Fus IDR-YFP) in the nucleus and at the plasma membrane. (C) Replacing the NC region with the HNRNPA1 IDR restored Gag’s ability to form discrete plasma membrane foci and the chimera was more strongly localized to the nucleus (RU5.Gag-HNRNPA1 IDR-YFP). (D) In the absence of the IDR, FUS was primarily diffuse but formed some aggregates. White arrows point out nuclear foci in (D) and (E). Replacing the FUS IDR with the NC IDR led to increased formation of foci in the cytoplasm and in the nucleus (FUS-NC- YFP). (F) Replacing the FUS IDR with MAp2p10 resulted in the formation of foci that localized primarily to the plasma membrane (FUS-MAp2p10-YFP). (G) In the absence of its IDR, HNRNPA1 was diffuse throughout the entire cell. (H) Replacing the HNRNPA1 IDR with NC induced the formation of cytoplasmic foci (HNRNPA1-NC-YFP). (I) Exchanging the HNRNPA1 IDR with MAp2p10 (HNRNPA1-MAp2p10-YFP) led to the formation of cytoplasmic foci with less targeting to the nucleus compared with the HNRNPA1-NC chimera. Nuclei were outlined in white dashed lines. Scale bar= 2 µm.

To determine whether this effect was specific for the FUS IDR, we used the IDR from HNRNPA1 as the donor. Replacing NC with the HNRNPA1 IDR resulted in an increase in the nuclear localization of Gag, and foci were formed primarily along the plasma membrane (Figure 9C). These data suggest that both the FUS and HNRNPA1 IDRs can functionally replace the NC sequence to restore the formation of Gag foci.

As a next step, we examined whether the Gag IDRs in MAp2p10 and NC could replace the FUS and HNRNPA1 IDRs. The full-length FUS-YFP protein forms foci in the nucleus and cytoplasm in QT6 cells, but is primarily nuclear in HeLa cells [73], suggesting a cell type difference (data not shown). When the FUS IDR was deleted, the protein was cytoplasmic and formed large aggregates rather than discrete foci (Figure 9D). The replacement of NC for the FUS IDR resulted in formation off discrete cytoplasmic and nuclear foci (Figure 9E). The substitution of the MAp2p10 IDR sequence for the FUS IDR led to plasma membrane-localized foci, like that of full-length Gag protein (compare Figure 1B with 9F). Deletion of the HNRNPA1 IDR caused the protein to localize in a diffuse pattern throughout the cell (Figure 9G). Replacing the HNRNPA1 IDR with NC or MAp2p10 restored formation of discrete foci in the cytoplasm and at the plasma membrane for both chimeric proteins (Figures 9H and 9I, respectively).

### Sensitivity of Gag BMCs to 1,6-Hexanediol *In vitro* and *In vivo*

Traditionally the aliphatic alcohol 1,6’-hexanediol has been used to study phase-contrasted BMCs because it dissolves droplets by disrupting weak hydrophobic protein-protein interactions [74]. For example, condensates formed by several nuclear proteins have been shown to be disrupted by 1,6’- hexanediol [75–79]. To determine whether RSV Gag foci would be disrupted by 1,6’-hexanediol, QT6 cells expressing WT Gag or nuclear-restricted Gag were treated with 10% 1,6’-hexanediol for 1 minute, and the paraspeckle protein NONO was used as a control (Figure 10A, panel a). With 1,6’-hexanediol treatment, a majority of NONO foci were disrupted, as expected, although some nuclear foci remained intact under these conditions, indicating that the global cellular architecture was not destroyed by drug treatment. When cells expressing WT Gag were treated with 1,6’-hexanediol, foci completely dissolved, and Gag became diffuse throughout the cell (panel b). Similarly, nuclear Gag foci formed by co-expression of NP214 dissolved with 1,6’- hexanediol treatment (panel c). The nuclear foci formed by Gag.L219A were mostly dissolved with 1,6’-hexanediol treatment, although some foci remained visible in the nucleus and along the plasma membrane (panel d). Although 1,6’-hexanediol is toxic and has nonspecific effects [74, 80], it should be noted that not all foci were dissolved, particularly for NONO, and cells remained intact during the short treatment period.

**Figure 10:**
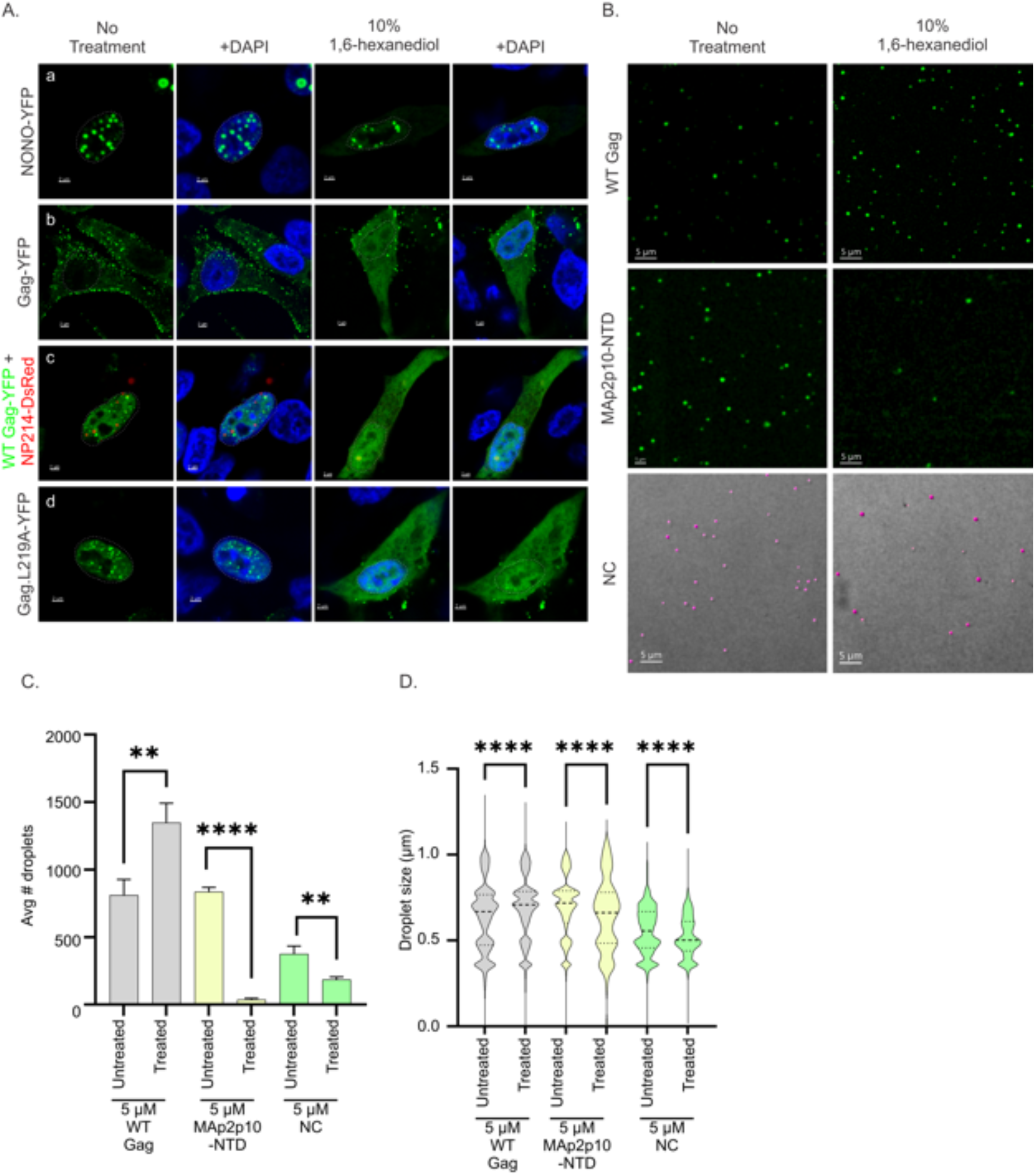
*In vivo* and *in vitro* perturnbation of BMCs with 1,6-hexanediol. A characteristic of BMCs is disruption by 1,6-hexandiol. QT6 cells expressing the indicated constructs (A) were incubated with 10% 1,6-hexanediol for 1 minute and in vitro droplets (B) were treated for 10 minutes. (A) panel a. NONO-YFP foci were disrupted by 1,6-hexanediol to a minor degree in QT6 cells. Panels b, c, and d. WT RU5.Gag-YFP and nuclear-restricted Gag foci became diffuse throughout the entire cell following treatment, indicating that Gag foci were sensitive to disruption in all cellular compartments. Scale bar= 2 µm. (B) The effect of 1,6-hexanediol on *in vitro* droplets was observed on recombinant WT Gag, MAp2p10-NTD, and NC proteins. 1,6- hexanediol (10%) was mixed with 5 µM of each protein in the presence of Ficoll-400 and NaCl, then incubated for 10 minutes before imaging. For better display of the NC droplets visualized using DIC, the image was overlaid with spheres generated by the spot function in Imaris (magenta). Scale bar= 5 µm. (C) and (D) 10 low-magnification fields were collected for each condition and subjected to the Imaris spot function to measure the number and size of droplets. In all cases, 1,6-hexanediol treatment induced statistically significant changes.

Next, we asked whether treatment with 10% 1,6’-hexanediol disrupted the formation of droplets *in vitro* (Figure 10B-D). Droplets were assembled using the method described in Figure 2, 10% 1,6-hexanediol was added for 10 minutes, and images were analyzed using the Imaris spot function. Droplets formed by WT Gag and MAp2p10-NTD were counted using the fluorescence channel and the DIC images were used for NC quantification. To better visualize the NC droplets for display purposes, the droplets were overlaid with the magenta spots in Imaris and enlarged (Figure 10B). Treatment with 1,6’-hexanediol had a significant effect on droplets for all three proteins, with a reduction in the number and size of the droplets formed by the isolated IDRs, MAp2p10 and NC. Interestingly, there was a paradoxical effect of 1,6’- hexanediol treatment on WT Gag droplets, which became larger and more numerous. The reason for this unexpected result could be due to effects other than weak hydrophobic forces holding the WT Gag complexes together *in vitro*. The different effect of hexanediol on *in vivo* and *in vitro* full-length Gag droplets could also be due to the drug’s effect on other cellular factors such as RNA, proteins, or lipids interacting with Gag.

## Discussion

It has been increasingly appreciated that BMCs play an important role the replication of many viruses, including influenza, SARS-CoV-2, HIV-1, and others, as reviewed elsewhere [81–84]. In our previous work, we observed the *in vivo* formation of RSV Gag foci in the nucleus, cytoplasm, and plasma membrane that colocalize with unspliced viral RNA and splicing factors [3, 4, 23]. Furthermore, we have observed that the Gag nuclear foci are dynamic and not merely aggregates [3, 22]. Based on the new data presented here, the RSV Gag polyprotein meets several criteria for BMCs that undergo LLPS [33, 34, 39, 50, 75, 85, 86], based on *in vivo* and *in vitro* assays. These properties include formation of foci that can be visualized using DIC microscopy (Fig. 1B-E); droplets with liquid-like properties that undergo fusion and fission in cells and *in vitro* (Figs. 4 and 5); complexes that exchange rapidly with the molecules in the surrounding environment, with a short half-time of recovery and prominent mobile fraction (Figs. 6 and 7; Tables 1-4); and disruption of foci upon treatment of cells with 1,6’-hexanediol (Fig. 10) [74, 87].

Viral and cellular proteins that undergo condensate formation typically contain IDRs that regulate affinity or specificity of binding and may also function as nucleic acid binding domains [88–90]. We identified two IDRs in Gag, one in the N-terminal region encompassing MAp2p10 and the other in the C-terminal NC domain. Both IDR sequences are sufficient to form BMCs *in vitro*, although MAp2p10 appears to play a more prominent role in Gag droplet formation.

Arguably, the most compelling evidence for their ability to provide the driving force in BMC formation is their ability to substitute functionally for the IDRs of FUS and HNRNPA1, and their ability to undergo location-specific condensation in optoDroplet assays. In addition, the need for an IDR function in Gag to properly form BMCs and direct it to the proper subcellular compartments is supported by the exchangeability of the FUS and HNRNPA1 IDRs for the NC IDR in Gag.

Interestingly, the Gag IDRs are located in regions that interact with cellular factors during the assembly process: MA contains an NLS that binds the karyopherins importin-11 and TNPO3 [16, 91]; p2 contains the late assembly domain [12] that binds to ESCRT proteins [92] and the NEDD4 ubiquitin ligase [93]; p10 contains an NES that interacts with the host export factor CRM1 [17, 24]; and NC contains a nuclear import signal that binds directly to importin-alpha to recruit importin-beta [16–19]. Additionally, MA has nonspecific nucleic binding activity, whereas NC possesses high affinity for the psi packaging sequence [2, 94–97]. We have previously shown that the Gag NC domain is required for the formation of nuclear Gag foci, suggesting that condensate formation depends on intermolecular protein-protein interactions and RNA binding [23]. Therefore, it is possible that the IDRs in RSV Gag play important functions in binding to host and viral factors that mediate transport of Gag from the nucleus to the plasma membrane during the process of creating virions. It has been proposed that IDRs adopt an ensemble of orientations that allow for promiscuous interactions with a variety of binding partners [98–102].

Many viral proteins contain IDRs that are used to hijack cellular processes [103, 104], including nucleocytoplasmic transport [105], which can help to explain how the MA region binds two different nuclear import factors, nucleic acids, and then serves as a plasma membrane-binding domain by interacting with acidic phospholipids [2, 106]. The same principle applies to the IDR in NC, which binds importin-alpha for nuclear import, interacts with nucleic acids nonspecifically, and binds the psi packaging sequence specifically [16, 24, 96, 107–111]. Further investigation will be required to sort out the roles of MA and NC with their respective nucleic acid binding activities in the formation of BMCs, and the complex interplay of other host factors that interact with each of them through their IDRs.

Our studies revealed some intriguing differences in the biophysical properties of WT Gag and the nuclear export mutant Gag.L219A. Imparted by a point mutation in the p10 NES of Gag and affecting a structurally important domain [18, 63, 112], Gag.L219A appears to have more dynamic properties and exhibits more liquid-like behavior compared to WT Gag, as evidenced by frequent fusion and fission events. The Gag.L219A protein accumulates in large nuclear foci, and it is defective in nucleocytoplasmic trafficking and virus particle formation. These observations lead us to propose that controlling phase transitions of Gag complexes are crucial for proper virus assembly. The fine-tuned multivalent interactions that promote Gag-Gag interactions allow the formation of compact, condensed foci that must maintain their integrity as they traffic through different compartments of the cell. However, dysregulation of these interactions can result in aberrant phase transitions [113–115], such as those induced by the L219A mutation, which blocks nuclear export, interferes with an important structural element in the polyprotein, and is defective in virus particle assembly. This idea is consistent with the concept that biological regulation of phase transition states is critical to maintaining functional condensates in cellular BMCs to prevent gels or fibrils from forming and causing disease [31, 38, 53, 54, 116–118].

We found it interesting that Gag foci remained similar in size and demonstrated fairly similar FRAP kinetics in the nucleus, cytoplasm, and plasma membrane. Further experiments will focus on identifying whether there are unique viral and cellular components of RSV BMCs in the nucleus, cytoplasm, and plasma membrane. Because most cellular proteins that form BMCs are located in a single subcellular compartment, RSV offers a unique opportunity to compare the biophysical properties of viral assembly complexes in different subcellular environments.

## Materials and Methods

### Plasmids

The plasmid pRU5.Gag.L219A-YFP, containing the RU5 RNA sequence followed by the *gag* coding region, was previously described [22]. pGag.L219A-CFP was previously described [4]. pRU5.Gag-YFP was cloned by replacing the L219A site in pRU5.Gag.L219A-YFP using restriction sites SacI and SpeI. pGag.L219A-YFP and pGag ΔNC-YFP were described previously [23]. pNP214-DsRed was described previously [19]. pYFP-PSP1alpha (referred to as YFP-PSP1 in this manuscript) was a kind gift was Dr. Angus Lamond, University of Dundee, United Kingdom [66]. pNONO-YFP (also known as p54/nrb) was previously described [3].

Plasmids pRU5.Gag-FUS IDR-YFP and pRU5.Gag-HNRNPA1 IDR-YFP were created through Gibson assembly [119]. Fragment 1 for both constructs was pRU5.Gag-YFP digested with NruI and ApaI to remove coding regions of CA and NC. Fragment 2 for both constructs consisted of CA coding region amplified from pRU5.Gag-YFP using primers: 5’ – CAC AAG ACT GGC TGA TAC GGT CAG GACC - 3’ and 5’ – CAT GGC CGC GGC TAT GCC TTG ATCC - 3’. For pRU5.Gag-FUS IDR-YFP, Fragment 3 consists of the *fus idr* coding region from pSNAP-FUS- IDR (a generous gift from Dr. Roy Parker, University of Colorado [118]) flanked by sequences overlapping the 3′ end of *ca* and the 5′ end of *yfp*, and was generated using primers: 5’ – GGA TCA AGG CAT AGC CGC GGC CAT GAT GGC CTC AAA CGA TTA TAC CCA ACA AG - 3’ and 5’ – GAC CGG CCG GTG GAT CCC GG CAC CAC TGC TGC GGT TGT AAC CAC - 3’.

For pRU5.Gag-HNRNPA1 IDR-YFP, Fragment 3 consists of the *hnrnpa1 idr* coding region from pET9d-HNRNPA1, which was a kind gift from Dr. Douglas Black (Addgene plasmid # 23026; http://n2t.net/addgene:23026; RRID: Addgene_23026) flanked by sequences overlapping the 3′ end of *ca* and the 5′ end of *yfp*, and was generated using primers: 5’ – GGA TCA AGG CAT AGC CGC GGC CATG ATG GCT AGT GCT TCA TCC AGC CAA AG - 3’ and 5’ – GAC CGG CCG GTG GAT CCC AA ATC TTC TGC CAC TGC CAT AGC TAC - 3’.

The construct pFUS ΔIDR-YFP was created by PCR amplifying the *fus* gene from pGST- TEV-FUS (a kind gift from Dr. Aaron Gitler; Addgene plasmid # 29629; http://n2t.net/addgene:29629; RRID: Addgene_29629) [120] with the addition of XhoI and BamHI sites using primers: 5’ – AGA TCT CGA GGC CAC CAT GTA TGA ACC CAG AGG TCG TGG AGGT – 3’ and 5’ - CGG TGG ATC CAA ATA CGG CCT CTC CCT GCG ATC CTG - 3’, and inserting the product into the pEYFP.N1 vector. Plasmid pFUS-MAp2p10-YFP and pFUS- NC-YFP were created by Gibson assembly using the same Fragments 1 and 2 for both constructs. Fragment 1 was pEYFP.N1 digested with ApaI. Fragment 2 was amplified *fus* minus the *idr* from pGST-TEV-FUS using primers: 5’ – TAT GAA CCC AGA GGT CGT GGA - 3’ and 5’ – GTG AAC AGC TCC TCG CCC TTG CTC AC TGA GAT ATC ACT ATAC GGC CTC TCC CTG CGA TCCT - 3’, which created an overlapping region to Fragment 1. Fragment 3 was designed from pFUS-MAp2p10-YFP to create overlapping regions to Fragments 1 and 2 using primers: 5’ – AGT CGA CGG TAC CGC GCC ACC ATG GAA GCC GTC ATA AAG GTG ATT TCG - 3’ and 5’ – TCC ACG ACC TCT GGG TTC ATA CAT GGC CAC CAC GGG CGG - 3’, which amplify *map2p10* from pRU5.Gag.YFP. Fragment 3 for pFUS-NC-YFP contained overlapping regions to Fragments 1 and 2 using primers: 5’ – AGT CGA CGG TAC CGC GCC ACC ATG GCA GTA GTC AAT AGA GAG AGG GAT GGA CA - 3’ and 5’ – TCC ACG ACC TCT GGG TTC ATA CGA GAC GGC AGG TGG CTC AGG - 3’, which amplify *nc* from pRU5.Gag.YFP.

To make pHNRNPA1ΔIDR-YFP, Gibson assembly was used. pEYFP.N1 was digested with AgeI and EcoRI and used as Fragment 1. pHNRNPA1ΔIDR-YFP Fragment 2 used primers 5’ – TAG CGC TAC CGG ACT CAG ATC TCG AGG CCA CCA TGT CTA AGT CAG AGT CTC CTA AAG AGCC - 3’ and 5’ – GGT GAA CAG CTC CTC GCC CTT GCT CAC GCC GCT ACC GCC CTC TTG CTT TGA CAG GGC TTT TCT AAC - 3’, which amplified *hnrnpa1* minus the *idr* sequence from pET9d-HNRNPA1.

Plasmids pHNRNPA1-MAp2p10-YFP and pHNRNPA1-NC-YFP were created through Gibson assembly where Fragment 1 was pEYFP.N1 digested with AgeI and EcoRI. To make pHNRNPA1-MAp2p10-YFP, Fragment 2 used primers 5’ – TAG CGC TAC CGG ACT CAG ATC TCG AGG CCA CCA TGT CTA AGT CAG AGT CTC CTA AAG AGC C - 3’ and 5’ – ATC ACC TTT ATG ACG GCT TCC TCT TGC TTT GAC AGG GCT TTTC - 3’, which amplified *hnrnpa1* minus the *idr* sequence from pET9d.HNRNPA1. Fragment 3 amplified *map2p10* from pRU5.Gag.YFP with overlapping sequences matching Fragments 2 and 3 with primers 5’ – GAA AAG CCC TGT CAA AGC AAG AGG AAG CCG TCA TAA AGG TGAT - 3’ and 5’ – GA ACA GCT CCT CGC CCT TGC TCA CGC CGC TAC CGCC ATAA GGA GGA GGA GGA GCC GA - 3’. To make pHNRNPA1-NC-YFP, Fragment 2 used primers 5’ – TAG CGC TAC CGG ACT CAG ATC TCG AGG CCA CCA TGT CTA AGT CAG AGT CTC CTA AAG AGC C - 3’ and 5’ – GTC CAT CCC TCT CTC TAT TGA CTA CTG CCT CT TGC TTT GAC AGG GCT TTT C - 3’ to amplify *hnrnpa1* minus the *idr* sequence from pET9d.HNRNPA1. Fragment 3 amplified *nc* from pRU5.Gag.YFP with overlapping sequences matching Fragments 2 and 3 with primers 5’ – GAA AAG CCC TGT CAA AGC AAG AGG CAG TAG TCA ATA GAG AGA GGG ATG GAC - 3’ and 5’ – GA ACA GCT CCT CGC CCT TGC TCA CGC CGC TAC CGCC CGA GAC GGC AGG TGG CTC AGG - 3’. pGag-SNAPTag was made by digesting pGag-CFP [23] with ApaI and NotI to remove the CFP, followed by PCR amplification of the SNAPTag from pSNAP-FUS-IDR [118] using primers that insert ApaI and NotI sites at the 5’ and 3’ ends, respectively: 5’ – ATC GGG GCC CGG GAT CCA CGA CAA AGA CTG CGA AAT GAA GCG CAC CACC - 3’ and 5’ – GCA TGC GGC CGC ATC GAT TTA ACC CAG CCC AGG CTT GCC CAG TCT - 3’.

The constructs encoding the Gag proteins used for purification [pET28(-His).Gag.ΔPR, pET28(-His).Gag.ΔSPΔNC, pET28(-His).MA.p2.p10.CA-NTD, pET28(-His).CA.NC, and pET28.MA) were described in [91]. pET28(-His).NC was made by digesting pET28a(- His).Gag.ΔPR with NdeI and HindIII. NC was amplified from pET28a(-His).Gag.ΔPR using appropriate primers (5’- TACG CATATG <u>GCA</u> GTA GTC AAT AGA GAG AGG GAT GGA CAA AC -3’ and 5’- GAT CAA GCT TTT ATT ATT ACG AGA CGG CAG GTG GCT CAG G -3’), and was inserted in between the NdeI and HindIII sites in the vector following digestion. To make pET28a(-His).Gag.L219A.ΔPR, pET28a(-His).Gag.ΔPR was digested with SpeI and ScaI to remove the wild-type p10, which was replaced with the mutant L219A region from pKoz.Gag.L219A.3h-YFP that was digested with the same enzymes [22]. pET24.CA was a gift from Rebecca Craven and was purified as described [121].

To create the constructs for the optoDroplet imaging, CRY2olig-mCherry, a gift from Chandra Tucker (Addgene plasmid # 60032; http://n2t.net/addgene:60032 ; RRID:Addgene_60032) [68]. To create FUS-IDR.CRY2olig-mCherry, FUS-IDR was amplified from pSNAP-FUS-IDR using primers: 5’- ACT GGC TAG CGC CAC CAT GGC CTC AAA CGA TTA TAC CCA ACA AGC -3’ and 5’- CAG TCT CGA GAC CAC TGC TGC GGT TGT AAC-3’, and inserted into the NheI/XhoI sites of CRY2olig-mCherry. All Gag sequences were amplified from pRU5.Gag.3h-YFP [22].CA.CRY2oligo-mCherry was created by amplifying CA with primers: 5’- ACT GGC TAG CGC CAC CAT GCC TGT AGT GAT TAA GAC AGA GGG ACC C- 3’ and 5’- CAG TCT CGA GCA TGG CCG CGG CTA TGC CTT G -3’. Because an XhoI site is present in Gag, the XhoI site in the original construct was replaced with an MluI site using Q5 site directed mutagenesis (New England Biolabs) using primers: 5’- GGA CTC AGA TAC GCG TGC CAC CAT GAA GAT GGA CAA AAA G - 3’ and 5’- GGT AGC GCT AGC GGA TCT - 3’.

All of the other Gag constructs were made by inserting PCR products amplified from pRU5- Gag.3h-YFP into the NheI and MluI sites of the modified CRY2olig-mCherry construct. RSV Gag.ΔPR-CRY2olig-mCherry was created by amplifying Gag.ΔPR using primers: 5’- ACT GGC TAG CGC CAC CAT GGA AGC CGT CAT AAA GGT GAT TTC G - 3’ and 5’- CAG TAC GCG TCG AGA CGG CAG GTG GCT CAG -3’. MAp2p10-CRY2olig-mCherry was generated by amplifying MAp2p10 using primers: 5’- ACT GGC TAG CGC CAC CAT GGA AGC CGT CAT AAA GGT GAT TTC G - 3’ and 5’- CAG TAC GCG TCA TGG CCA CCA CGG GCG G-3’. NC- CRY2oligo-mCherry was created by amplifying NC using primers: 5’- ACT GGC TAG CGC CAC CAT GGC AGT AGT CAA TAG AGA GAG GGA TGG ACA - 3’ and 5’- CAG TAC GCG TCG AGA CGG CAG GTG GCT CAG -3’. Expression of these constructs in HEK293T cells was verified by Western blot alongside CRY2oligo-mCherry alone, which served as a negative control. Plasmid sequences were all confirmed by DNA sequencing (Eurofins).

### Cell Lines, Transfection, Fixation

Chemically transformed quail fibroblast QT6 cells [122] were maintained as described [123]. HEK293T cells obtained from ATCC (CRL-3216) were grown in Dulbecco’s Modified Eagle Medium (DMEM), 10% FBS, 1% sodium pyruvate, penicillin/streptomycin and fungizone at 37°C. Cells were seeded onto coverslips for fixed cells or on MatTek dishes for live-cell imaging.

QT6 Cells were transfected via the calcium phosphate method [124] using the following amounts of the indicated plasmid: pGag-SNAPTag (500 ng), pNP214-DsRed (1 µg), pRU5.Gag-YFP (500 ng), pGag.L219A.YFP/CFP (1.5 µg), pGag ΔNC-YFP (1 µg), pYFP-PSP1 (100 ng), pNONO-YFP (100 ng), pFUS ΔIDR-YFP (500 ng), pFUS-MAp2p10-YFP (500 ng), pFUS-NC- YFP (500 ng), pHNRNPA1 ΔIDR-YFP (500 ng), pHNRNPA1-MAp2p10-YFP (500 ng), pHNRNPA1-NC-YFP (500 ng), pRU5.Gag-FUS IDR-YFP (500 ng), and pRU5.Gag-HNRNPA1 IDR-YFP (500 ng). Cells containing SNAPTag fusion proteins were labeled with either SNAPTag ligand JF549 or JF646 (a kind gift from Luke Lavis, Janelia Research Campus [125]) and incubated for another hour at 37°C. For fixed cell experiments, cells were fixed in either 3.7% paraformaldehyde (PFA) in PHEM buffer [3.6% piperazine-*N,N*=-bis(2-ethanesulfonic acid) (PIPES), 1.3% HEPES, 0.76% EGTA, 0.198% MgSO4, pH 7.0) [126] for 20 minutes or 3.7% formaldehyde in PBS for 10 minutes, washed with PBS, 4’,6-diamidino-2-phenylindole (DAPI) stained, and mounted in either antifade reagent (Invitrogen) or ProLong Diamond (Life Technologies).

For visualizing Gag foci in fixed QT6 cells (Figure 1), imaging was performed using a Leica AOBS SP8 confocal microscope with a 63x/1.4 oil objective. DAPI was excited with the 405 nm UV laser at 10% laser power and detected using a photomultiplier tube detector (PMT). The following were excited with the WLL at the specified laser lines and detected with a hybrid detector (HyD): Gag-SNAPTag-JF646 at 646 nm, NP214-dsRed at 558 nm, and Gag.L219A- CFP at 470 nm. Differential interference contrast (DIC) or brightfield (BF) were excited at 488 nm and detected with a PMT.

For live cell imaging of optoDroplets, HEK293T cells were seeded at a density of 1x10^6^ cells/dish in 35-mm glass-bottom dishes (MatTek) and were allowed to grow to ∼80% confluency. Cells were then transfected for ∼16 hours with 0.5 µg of DNA, using jetOptimus transfection reagent (Polyplus). Dishes were placed in a dark incubator post-transfection and were protected from blue light exposure from this point onwards. One hour prior to imaging, the live cell chamber was prepared and allowed to equilibrate to 37°C/5% CO2; 30 mins prior to imaging, media was removed from the MatTek dishes and replaced with Imaging Media (Fluorobrite DMEM High Glucose, without glutamine (Gibco) + 10% FBS + 1% sodium pyruvate). A Leica SP8 FALCON microscope equipped with an incubated stage at 37°C/5% CO2 was used to collect images. Cells were imaged in the mCherry channel (584 nm) at 5 second intervals for ∼30 seconds, then were exposed to blue light (488 nm; 0.02% laser power) and images were captured at 5 second intervals for 300 seconds. All live cell imaging was performed using a 63X/1.2 water objective at 3X magnification. Images and timelapse videos were prepared using Imaris software.

### Leptomycin B (LMB) Treatment

QT6 cells were transfected with pGag-SNAPTag for 14 hours, then 1 mL of QT6 medium containing 10 ng/mL LMB was added and incubation continued for 1 hour at 37°C. The medium was swapped for 1 mL of QT6 medium containing 10 ng/mL LMB and 1 µM SNAPTag ligand JF646 (a kind gift from Luke Lavis, Janelia Research Campus [125]) and incubated for another hour for a total treatment time of 2 hours. Cells were then fixed in 3.7% formaldehyde in PBS for 10 minutes, washed with PBS, DAPI stained, and mounted with ProLong Diamond (Life Technologies) before being imaged on a Leica AOBS SP8 confocal microscope with a 63x/1.4 oil objective. DAPI was excited with the 405 nm UV laser at 10% laser power and detected using a photomultiplier tube detector (PMT), and JF646 was excited using a white light laser (WLL) with a laser line excitation of 646 nm and detected using a HyD.

### *In Vitro* Droplet Formation

Recombinant RSV WT and mutant Gag proteins were expressed from pET28 (-His) vector in *E. coli* BL21(DE3) pRIL cells, purified through a HPLC SEC-MALS column, and stored in standard buffer (300 to 500 mM NaCl, 10 mM HEPES, pH 7.5, 0.1 mM EDTA; 0.1 mM TCEP; 0.01 mM ZnCl2). Nucleic acid contamination was assessed by measuring absorbance at 280 nm and 260 nm, with ratios indicating the absence of nucleic acid contamination, as described previously [24]. Purified proteins were labelled using the Alexa Fluor 488 Microscale Protein Labeling Kit (Thermo Fisher Scientific, #A30006) according to manufacturer’s protocol. Labelled protein was incubated with unlabeled protein (at a ratio of either 1:10 or 50:50) [127] to reach a final concentration of 5, 10, or 20 µM in buffer containing 20 mM HEPES, pH 7.5-8.0, 150-250 mM NaCl, and 86-150 mg/ml Ficoll-400 (used as a crowding agent), modified from: [62].

Immediately after mixing, 6-10 µl of the protein solution was put on a coverslip and a slide was placed on top. Coverslips were sealed with clear nail polish, then imaged after 10 minutes using the Leica AOBS SP8 confocal microscope with a 63x/1.4 oil objective. TA WLL laser line excitation of 493 nm was used in conjunction with a hybrid detector to detect the 488 fluorescent label, and DIC was used to get phase contrasted images of the droplets.

Droplet numbers and sizes were obtained from low magnification images using the Imaris spot function, and only spherical single droplets were counted. Prior to analysis, background was removed using a Gaussian filter. For all conditions, the fluorescence channel was used to count droplets except in the case of NC and CA. Of note, proteins were fluorescently labelled with Alexa Fluor 488 (see above) by binding to primary amines, of which NC and CA have only 7 and 9, respectively.

Statistics were determined by one-way ANOVA using GraphPad Prism (GraphPad Software, Inc.). The phase diagram of WT Gag (RSV Gag ΔPR) was determined using the following method. Ficoll-400 was dissolved to yield a final concentration of 10% w/v in a solution containing 20 mM HEPES-OH, pH 7.5, and varying NaCl concentration (as indicated) and Gag protein was added with gentle mixing. Immediately, 6 µl was deposited on an 18 mm square glass coverslip and a slide was placed on top of the coverslip. Slides were incubated at room temperature for 5 minutes, then sealed with clear nail polish and placed in a laminar flow hood to dry for an additional 5 minutes. A Nikon CSU-X1 Spinning Disk Field Scanning Confocal microscope was utilized to collect images of 10 fields per condition at 100X magnification. Condensate number was quantified in Imaris (Bitplane) using the spot function, and heat map generated using GraphPad Prism (GraphPad Software, Inc.).

### Live Cell Imaging

Living cells, expressing either Gag-SNAPTag or Gag.L219A-YFP, were imaged on a Leica AOBS SP8 confocal microscope with a 63x/1.2 water objective and a live cell chamber at 37°C with 5% CO2. QT6 cells expressing Gag.SNAPTag were labelled with 100 nM of Janelia Fluor 549 (JF549) ligand [125] in imaging media [clear Dulbecco’s Modified Eagle Medium (DMEM) with 4.0 mM L-glutamine and 4.5 mg/liter glucose (Hyclone), supplemented with 5% fetal bovine serum, 9% tryptose phosphate broth, and 1% chicken serum] for 1 hour at 37°C. Media was exchanged with fresh imaging media and then the cells were imaged for 15 minutes, with one frame imaged every minute. Cells expressing Gag.L219A-YFP were imaged for 16 minutes at one frame per minute. YFP was imaged using the WLL with a laser line excitation of 514 nm using a hybrid detector.

For time lapse imaging of Gag droplet fusion *in vitro*, solutions containing either unlabeled 20 µM Gag.ΔPR or Gag.L219A were mixed with crowding agent as described above, added to a coverslip, covered with a slide, sealed with nail polish, and imaged using BF or DIC microscopy using a WLL at 488 nm, and detected using a PMT, capturing a frame every 10 seconds.

### Fluorescence Recovery After Photobleaching (FRAP)

QT6 cells were seeded on glass-bottom 35 mm dishes (MatTek Corporation) and incubated overnight to allow them to adhere. The following day, cells were transfected for 16 hours, and the media was exchanged with imaging media. Cells were imaged using the FRAP module on the Leica AOBS SP8 confocal microscope 63x/1.2 water objective at 7.5x zoom with the pinhole set to 2 airy units. Three ROIs were selected: (1) the focus to be bleached, (2) the entire nucleus/cell to monitor and correct for whole cell bleaching, (3) outside of the cell for background subtraction. Ten frames were imaged of pre-bleached cells every 0.08 seconds using the Argon laser line at 514 nm set to 2% power. The laser was used at 100% power to bleach samples, for 15 frames every 0.08 seconds. Post-bleach recovery was imaged with laser line at 514 nm and 2% power over different iterations: 30 frames with 1 frame every 0.076 seconds, then 15 frames imaged every 1 second, and finally, more frames every 5 seconds. For analysis purposes, later timepoints were excluded for some samples such that all conditions had the same number of timepoints (end times ranged from 18 seconds to 35 seconds). Foci were bleached to at least 40% of the starting fluorescence intensity. To calculate the mobile fractions for each time point, the web application easyFRAP-web (https://easyfrap.vmenet.upatras.gr) was used [128]. Data was full scale normalized. GraphPad Prism 5 (GraphPad Software, Inc.) was used to generate plots of the mobile fractions over time.

### Transfections of Plasmids Expressing Gag-FUS and Gag-HNRNPA1 Chimeras

Following a 16-hour transfection, QT6 cells were fixed using 3.7% formaldehyde in PBS, washed with PBS, DAPI stained, and mounted in Prolong Diamond (Invitrogen). Slides were imaged on Leica AOBS SP8 confocal microscope with a 63x/1.4 oil objective. DAPI was excited with the 405 nm UV laser at 10% laser power using a PMT, and YFP imaged using the WLL with a laser line excitation of 514 nm using a hybrid detector.

### 1,6’-Hexanediol Treatment

Before seeding, coverslips were treated with a 2% w/v gelatin solution (DIFCO). The 2% solution was first autoclaved for 30 minutes. Coverslips were washed in 70% ethanol and allowed to dry. The coverslips were incubated in the gelatin solution for at least 10 seconds then propped up and allowed to dry for at least 2 hours. Following a 16-hour transfection, QT6 cells were either left untreated or were treated with 10% w/v 1,6’-hexanediol (Tokyo Chemical Industry Company, LTD) in media for 1 minute. Cells were then immediately fixed using 3.7% paraformaldehyde (PFA) in PHEM buffer (see above) for 20 minutes, washed with PBS, (DAPI) stained, and mounted in antifade reagent (Invitrogen). Slides were imaged on Leica AOBS SP8 confocal microscope with a 63x/1.4 oil objective. DAPI was excited with the 405 nm UV laser at 20% laser power using a PMT, and YFP was imaged using the WLL with a laser line excitation of 514 nm using a hybrid detector. DsRed was imaged using the WLL with a laser line excitation of 558 nm using a hybrid detector.

For hexanediol treatment of *in vitro* droplets, 50% w/v hexanediol was added to a final concentration of 10%, incubated with 5 µM of each Gag protein for 10 minutes, and imaged as previously described. Droplet numbers and sizes were obtained from low magnification images using the Imaris spot function as described above. Statistical analysis was performed to compare treated and untreated samples using an unpaired student’s t-test in GraphPad Prism (GraphPad Software, Inc.).

## Acknowledgements

We would like to acknowledge those who aided in this work. We are grateful for Angus Lamond (University of Dundee), Roy Parker (University of Colorado Boulder), and Rebecca Craven (Penn State College of Medicine), who generously provided plasmids. Luke Lavis (HHMI Janelia Research Campus) kindly provided the SNAPTag JF549 and JF646 ligands. We thank Alan Cochrane (University of Toronto), Andrew Mouland (McGill University), and Jordan Chang (Penn State College of Medicine) for critical discussions. This work was supported by a grant from the National Institutes of Health, R01 GM139392 (L.J.P.) and a Summer Bridge Award from the Penn State College of Medicine (L.J.P).

Microscopy images and were generated and processed in the Penn State College of Medicine Advanced Light Microscopy Core (RRID: SCR_022526). The Advanced Light Microscopy Core services and instruments used in this project were funded, in part, by the Pennsylvania State University College of Medicine via the Office of the Vice Dean of Research and Graduate Students and the Pennsylvania Department of Health using Tobacco Settlement Funds (CURE). The content is solely the responsibility of the authors and does not necessarily represent the views of the University or College of Medicine. The Pennsylvania Department of Health specifically disclaims responsibility for any analyses, interpretations or conclusions.

---

**Supplemental Table 1: *in vitro* droplet size comparison.**

**Supplemental Table 2: *in vitro* droplet number comparison.**

**Supplemental Movie 1**: To determine whether WT Gag droplets formed *in vitro* had liquid-like properties, 20 µM of unlabeled WT Gag protein mixed with crowding agent was imaged under brightfield every 10 seconds. A small droplet is seen fusing into a large droplet (black circle). Scale bar = 1 µm.

**Supplemental Movie 2**: To determine whether Gag.L219A *in vitro* droplets had liquid-like properties, 20 µM of unlabeled protein mixed with crowding agent was imaged by DIC every 10 seconds. Several examples of droplet fusion are outlined in black circles. Scale bar = 1 µm.

**Supplemental Movies 3a, 3b, and 3c: WT Gag *in vivo* fusion.** To examine fusion and fission of WT Gag in different subcellular compartments, a living QT6 cell expressing Gag-SNAPTag was imaged at 1 frame/minute. For better visualization of the foci in each subcellular compartment, images were adjusted in Imaris for the (a) nucleus, (b) cytoplasm, and (c) plasma membrane. Foci of interest in each movie were circled in white (nucleus), yellow (cytoplasm), or magenta (plasma membrane). Scale bar= 1 µm.

**Supplemental Movie 4: Nuclear restricted Gag *in vivo* fusion.** QT6 cells expressing the Gag.L219A-YFP NES mutant were imaged at 1 frame/minute. Several examples of fusing droplets were outlined in black circles. Scale bar= 1 µm.

**Supplemental Movie 5: CRY2oligo-mCherry time course.** HEK293T cells expressing CRY2oligo-mCherry were imaged every 5 seconds for 300 seconds. At 30 seconds, cluster formation was induced by the addition of blue light. CRY2oligo-mCherry formed few clusters. Scale bar = 2 µm.

**Supplemental Movie 6: FUS.IDR.CRY2oligo-mCherry time course**. Cells were imaged every 5 seconds for 300 seconds to detect mCherry fluorescence. FUS.IDR.CRY2oligo-mCherry underwent marked clustering in HEK293T cells when illuminated with blue light at 30 seconds. Scale bar = 2 µm.

**Supplemental Movie 7: Gag.CRY2oligo-mCherry time course.** HEK293T cells expressing Gag.CRY2oligo-mCherry were imaged every 5 seconds for 300 seconds to detect mCherry fluorescence. Cells were illuminated with blue light after 30 seconds of imaging, inducing the Gag.CRY2oligo-mCherry protein to form clusters at the plasma membrane. Scale bar = 2 µm.

**Supplemental Movie 8: MAp2p10.CRY2oligo-mCherry time course.** HEK293T cells expressing MAp2p10.CRY2oligo-mCherry were imaged every 5 seconds for 300 seconds to detect mCherry fluorescence. Although a few cytoplasmic foci are observed in the absence of blue light, illumination of the cells with blue light at 30 seconds caused MAp2p10.CRY2oligo-mCherry to cluster at the plasma membrane in HEK293T cells. Scale bar = 2 µm.

**Supplemental Movie 9: NC.CRY2oligo-mCherry time course.** When illuminated with blue light, NC.CRY2oligo-mCherry formed numerous clusters in the nuclei of HEK293T cells. Cells were imaged every 5 seconds for 300 seconds to detect mCherry fluorescence. Blue light was added 30 seconds into imaging session. Scale bar = 2 µm.

**Supplemental Movie 10: CA.CRY2oligo-mCherry time course.** The CA sequence does not contain a predicted IDR. When HEK293T cells expressing CA.CRY2oligo-mCherry were stimulated with blue light, the protein did not form clusters. Cells were imaged every 5 seconds for 300 seconds. Blue light was added after 30 seconds of imaging. Scale bar = 2 µm.

## Footnotes

1 Abbreviations: BMCs, Biomolecular condensates; RSV, Rous sarcoma virus; MA, matrix; CA, capsid; NC, nucleocapsid; PR, protease; IDRs, intrinsically disordered regions; vRNP, viral ribonucleoprotein complex; HIV-1; human immunodeficiency virus type 1; LMB, leptomycin B; NLS, nuclear localization signal; NES, nuclear export signal; DIC, differential interference contrast; FRAP, fluorescence recovery after photobleaching; NTD, N-terminal domain

